# Molecular basis of convergent evolution of ACE2 receptor utilization among HKU5 coronaviruses

**DOI:** 10.1101/2024.08.28.608351

**Authors:** Young-Jun Park, Chen Liu, Jimin Lee, Jack T Brown, Chen-Bao Ma, Peng Liu, Qing Xiong, Cameron Stewart, Amin Addetia, Caroline J. Craig, M. Alejandra Tortorici, Abeer Alshukari, Tyler Starr, Huan Yan, David Veesler

**Author notes:** These authors contributed equally to the work (Young-Jun Park, Chen Liu).

## Abstract

DPP4 was considered a canonical receptor for merbecoviruses until the recent discovery of African bat-borne MERS-related coronaviruses using ACE2. The extent and diversity with which merbecoviruses engage ACE2 and their receptor species tropism remain unknown. Here, we reveal that HKU5 enters host cells utilizing *Pipistrellus abramus* (P.abr) and several non-bat mammalian ACE2s through a binding mode distinct from that of any other known ACE2-using coronaviruses. These results show that several merbecovirus clades independently evolved ACE2 utilization, which appears to be a broadly shared property among these pathogens, through an extraordinary diversity of ACE2 recognition modes. We show that MERS-CoV and HKU5 have markedly distinct antigenicity, due to extensive genetic divergence, and identified several HKU5 inhibitors, including two clinical compounds. Our findings profoundly alter our understanding of coronavirus evolution and pave the way for developing countermeasures against viruses poised for human emergence.

## Introduction

Coronaviruses comprise a large group of pathogens circulating in mammalian and avian hosts. 229E, NL63, HKU1, and OC43 are endemic in the human population. SARS-CoV-1 caused an epidemic in 2002-2004 with an approximately 8% fatality rate^1,2^. MERS-CoV is highly lethal, with a 36% case fatality rate, and has been circulating sporadically since its discovery in 2012^3–5^ whereas SARS-CoV-2 is responsible for the COVID-19 pandemic^6,7^ with recurrent waves sweeping through the population^8,9^. SARS-CoV-1 and SARS-CoV-2 (*Sarbecovirus* subgenus) originated in bats^10–14^ and might have been transmitted to humans through intermediate hosts^15–17^. MERS-CoV (*Merbecovirus* subgenus) possibly originated in bats although spillovers occurred repeatedly from camels^5,18–22^, emphasizing the zoonotic threat of these coronaviruses.

The spike glycoprotein (S) is anchored in the viral membrane and required for cellular entry. S comprises an N-terminal S_1_ subunit (receptor-binding) and a C-terminal S_2_ subunit (membrane fusion)^23–25^ and is a main target of neutralizing antibodies, a correlate of protection against coronaviruses^26–31^. Host protease-mediated S cleavage occurs at the S_1_/S_2_ junction for many coronaviruses, yielding non-covalently linked S_1_ and S_2_ subunits in the prefusion S trimer, and at the S_2_’ site (adjacent to the fusion peptide) for all coronaviruses, leading to large-scale fusogenic conformational changes^32,33^. Coronavirus invasion of target cells thus requires the concerted action of receptor-binding and proteolytic S processing to lead to productive infection and both factors modulate the ability of coronaviruses to cross-species barrier.

Our ability to prepare for zoonotic spillovers is undermined by the knowledge gap between metagenomics discovery of a large number of coronaviruses and our understanding of the likelihood of their spillover to the human population. This is compounded by the ability of coronaviruses to undergo extensive antigenic changes and recombinations which can alter host receptor tropism through ortholog adaptation or receptor switch. For instance, N501Y-harboring SARS-CoV-2 variants acquired the ability to use mouse ACE2^34^ which was further enhanced with the addition of the Q493R mutation^8,35^. Furthermore, a recombination event in the merbecovirus S glycoprotein led to an unexpected receptor switch with MERS-CoV and HKU4 utilizing DPP4^36–38^ whereas a few African bat-borne merbecoviruses use ACE2^39,40^. However, the extent of and diversity with which merbecoviruses engage ACE2 and their host receptor tropism remain poorly understood.

Here, we show that the HKU5 clade of merbecoviruses, which was first described in 2006 in *Pipistrellus abramus*^41^ (P.abr), can unexpectedly use ACE2 from P.abr and from several Artiodactyl species for cell entry. We determined a cryo-electron microscopy structure of the HKU5 receptor-binding domain (RBD) complexed with P.abr ACE2, revealing a binding mode distinct from that of any other known ACE2-using coronaviruses, and identified the molecular determinants of receptor species tropism. We show that although MERS-CoV infection-elicited polyclonal plasma antibodies exhibit very weak to no HKU5 cross-neutralization, several broad-spectrum coronavirus inhibitors (including two clinical compounds) block HKU5 propagation in human cells. Our findings unveil that the HKU5 clade of merbecovirus independently evolved ACE2 utilization, underscoring the extraordinary diversity of ACE2 recognition modes among coronaviruses, and paving the way for developing HKU5 countermeasures against a clade of pathogen poised for human spillover.

## Results

### HKU5 utilizes ACE2 as entry receptor

Merbecoviruses comprise a large diversity of pathogens found in humans, bats, camels, pangolins and hedgehogs^1,2,5,18–22,37,42^. Phylogenetic classification based on their S receptor-binding domain (RBD) amino acid sequences delineates at least six merbecovirus clades **(Fig 1A)**, many of which do not have a confirmed host receptor. To investigate the receptor usage of the HKU5 merbecovirus clade, we evaluated binding of a dimeric HKU5-19s RBD N-terminally fused to a human Fc fragment (RBD-hFc) to the surface of HEK293T cells transiently transfected with a panel of 117 mammalian ACE2 orthologs across the phylogeny (**Fig 1B**). Although we only detected binding to P.abr ACE2 among the panel of 62 bat orthologs tested, the HKU5-19s RBD also bound to several non-bat mammalian ACE2s, including *Bos taurus* (B.tau), *Ovis aries* (O.ari), *Bos mutus* (B.mut), *Odocoileus virginianus texanus* (O.vir), *Mustela erminea* (M.erm), and *Bubalus bubalis* (B.bub) ACE2s (**Fig 1C-D and Fig S1A-B**). Furthermore, we observed efficient entry of vesicular stomatitis virus (VSV) pseudotyped with HKU5-19s S into HEK293T cells upon transfection with P.abr, B.tau, O.ari, B.mut, O.vir, M.erm, B.bub ACE2s as well as with the R.nor ACE2 (**Fig 1C-D and Fig S1C-D**). The observation of pseudovirus entry mediated by R.nor ACE2 for which no HKU5-19s RBD binding was detected likely results from avidity due to multivalent presentation of S trimers on pseudoviruses along with the use of trypsin to enhance pseudovirus entry^39,43,44^. Caco-2 cells stably expressing P.abr ACE2 (Caco-2/P.abr ACE2) enabled robust propagation of both a propagation-competent recombinant VSV encoding HKU5-19s instead of VSV-G (psVSV-HKU5-19s) and authentic HKU5-1, lending further support to the P.abr ACE2 receptor function (**Fig 1E-F**). Addition of exogenous trypsin promoted detectable (albeit weak) replication in human Caco-2 cells of authentic HKU5-1^45^, in line with prior data^46^. These findings indicate that HKU5 utilizes its native bat host ACE2 as entry receptor, concurring with a recent report^47^, along with several non-bat mammalian ACE2s from the Artiodactyl, Carnivore and Rodent orders.

**Figure 1.**
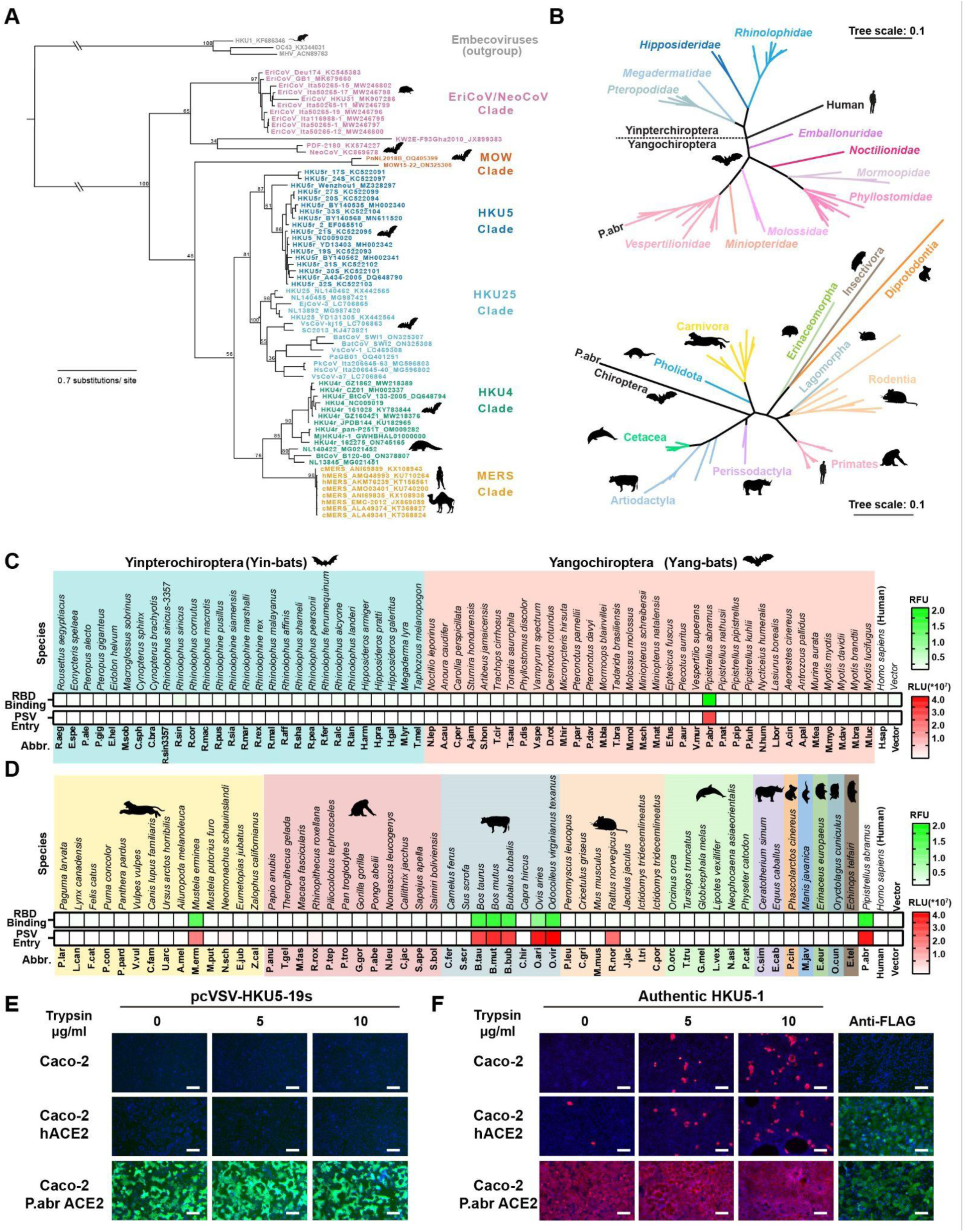
Identification of several mammalian ACE2s as functional HKU5 entry receptors. **(A)** Merbecovirus RBD phylogenetic tree based on amino acid sequences defining 6 clades. c/hMERS: camel/human MERS-CoV isolates. Each merbecovirus is listed along with its GenBank ID. The animal symbols represent the hosts in which viruses from a given clade have been detected. **(B)** Phylogenetic trees of bat (top) or non-bat (bottom) mammalian ACE2 orthologs based on amino acid sequences, with genera and orders indicated for the bat or non-bat mammalian species, respectively. **(C-D)** Binding of the HKU5-19s RBD-hFc to and entry of HKU5-19s S VSV pseudovirus into HEK293T cells transiently transfected with the indicated bat (C) or non-bat (D) mammalian ACE2 orthologs. Abbr: abbreviations used for species names. **(E-F)** Propagation of psVSV-HKU5-19s (E) and authentic HKU5-1 (F) in wildtype human Caco-2 cells or Caco-2 cells with stable expression of either human or P.abr ACE2. The propagation of psVSV-HKU5-19s was examined by the expression of the GFP reporter gene at 24 hours post-infection (hpi). The propagation of authentic HKU5-1 was detected by immunofluorescence with an anti-HKU5 N antibody at 24 hpi. The trypsin concentration used is indicated. ACE2 expression was confirmed by immunofluorescence using C-terminal-fused FLAG tags. Scale bars: 200 μm. Mean values are shown in C and D with n=3 biological replicates.

### Molecular basis of HKU5 recognition of ACE2

To understand HKU5 engagement of its host receptor, we characterized the HKU5-19s RBD in complex with a P.abr ACE2 ectodomain construct comprising the peptidase and the dimerization domains using cryoEM. Data processing revealed a predominance of a monomeric form of ACE2 (accounting for 99% of particles in selected 2D class averages despite the presence of the native dimerization domain) which led us to determine an asymmetric 3D reconstruction focused on the ACE2 peptidase domain and bound RBD at 3.1 Å resolution (**Fig 2, Fig S2 and Table 1**). The HKU5 RBD binds to the P.abr ACE2 peptidase domain and buries a surface of ∼950 Å^2^ from each of the two binding partners through interactions dominated by hydrogen bonding and shape complementarity.

**Figure 2.**
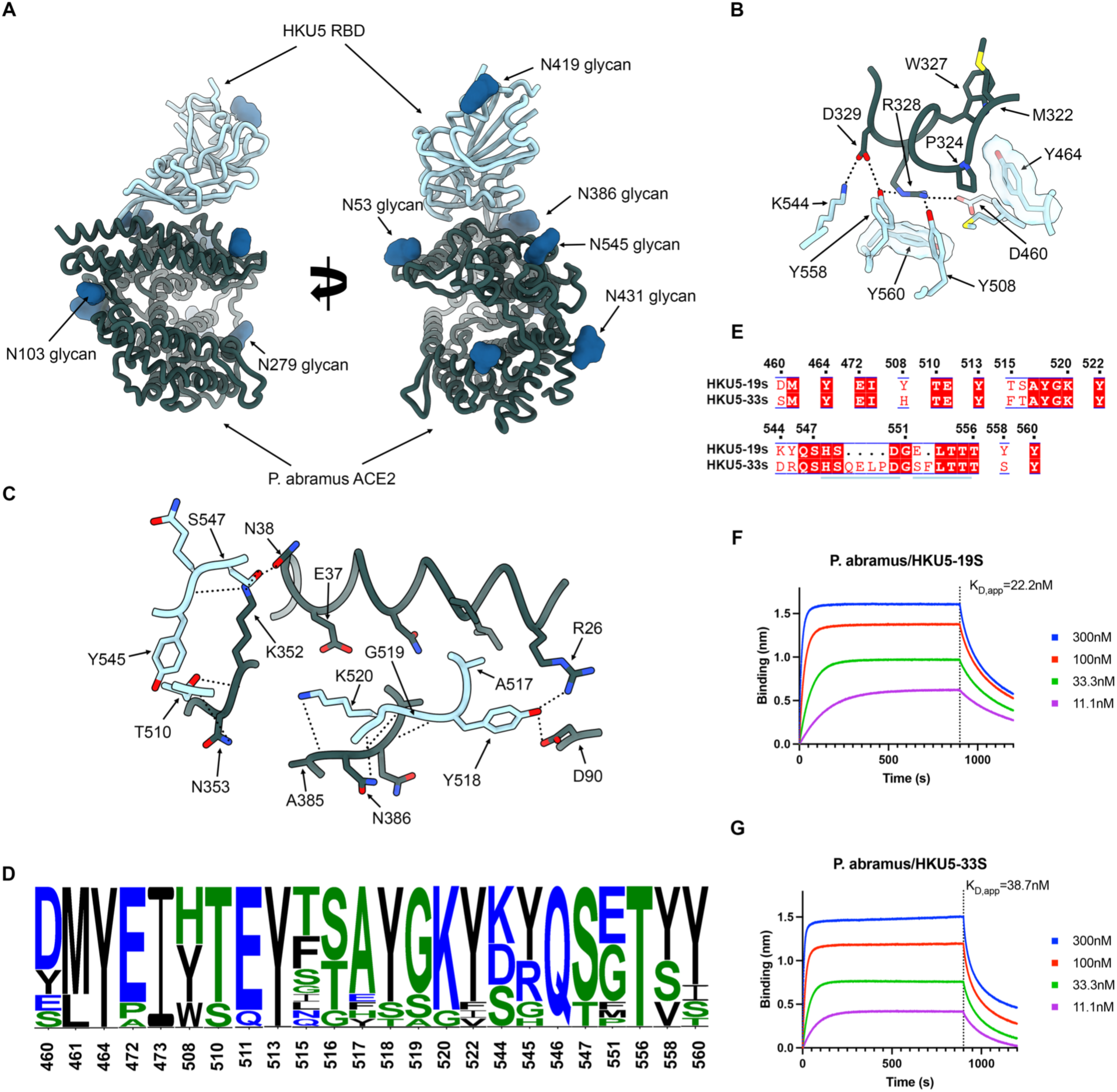
Molecular basis of HKU5 recognition of the P. abramus ACE2 receptor. **(A)** Ribbon diagrams in two orthogonal orientations of the cryoEM structure of the HKU5-19s RBD (light blue) bound to the P. abramus ACE2 peptidase domain (green) at 3.1Å resolution. **(B-C)** Zoomed-in views of the interface highlighting key interactions between the HKU5-19s RBD and P. abramus ACE2. Selected polar interactions are shown as black dotted lines. **(D)** Polymorphism of ACE2-interacting residues (RBM) among HKU5 isolates shown as logoplot. **(E)** RBM amino acid sequence alignment of the HKU5-19s and HKU5-33s isolates. Conserved residues are rendered with a white font over a red background whereas non-conserved residues are rendered with a red font on a white background. The residue numbering corresponds to HKU5-19s. HKU5-33s insertions are shown as black dots. The blue lines indicate residues outside the RBM shown for visualization purposes around the HKU5-33s insertions. **(F-G)** Biolayer interferometry analysis of the P.abr ACE2 ectodomain binding to the HKU5-19s and HKU5-33s RBDs immobilized on biolayer interferometry streptavidin (SA) biosensors. Binding avidities were determined by steady state kinetics and are reported as apparent affinities (K_D_, app) due to avidity.

**Table 1.** CryoEM data collection and refinement statistics.

|  |  |
| --- | --- |
|  | HKU5 RBD – P.abr ACE2 |
|  | PDB 9D32 |
|  | EMD 46512 |
| **Data collection and processing** |  |
| Magnification | 105,000 |
| Voltage (kV) | 300 |
| Electron exposure (e<sup>-</sup>/Å<sup>2</sup>) | 60 |
| Defocus range ($\mu\text{m}$) | -0.2 - -3.0 |
| Pixel size ($\text{\AA}$) | 0.843 |
| Symmetry imposed | C1 |
| Initial particle images (no.) | 3,159,512 |
| Final particle images (no.) | 550,684 |
| Map resolution ($\text{\AA}$) | 3.1 |
| FSC threshold | 0.143 |

**Table 1. CryoEM data collection and refinement statistics.**
|  |  |
| --- | --- |
| Model resolution ($\text{\AA}$) | 3.2 |
| FSC threshold | 0.5 |
| Map sharpening $B$ factor ($\text{\AA}^2$) | -135 |
| Model composition |  |
| Non-hydrogen atoms | 6140 |
| Protein residues | 778 |
| Ligands | 9 |
| $B$ factors ($\text{\AA}^2$) | |
| Protein | 15.0 |
| Ligand | 18.5 |
| R.m.s. deviations |  |
| Bond lengths ($\text{\AA}$) | 0.011 |
| Bond angles ($^\circ$) | 1.360 |
| Validation |  |
| MolProbity score | 1.39 |
| Clashscore | 3.55 |
| Poor rotamers (%) | 1.41 |
| Ramachandran plot |  |
| Favored (%) | 97.28 |
| Allowed (%) | 2.59 |
| Disallowed (%) | 0.13 |
| EMRinger score | 4.11 |

The HKU5 receptor-binding motif (RBM) folds as a compact four-stranded antiparallel β-sheet flanked by short α-helices on both sides and forms two consecutive insertions in the five-stranded antiparallel RBD core β-sheet (**Fig 2A**). Specifically, HKU5 residues D460, M461, Y464, E472, I473, Y508, T510, E511, Y513, T515, S516, A517, Y518, G519, K520, Y522, K544, Y545, Q546, S547, G551, T556, Y558 and Y560 form the RBM which interacts with ACE2 residues R26, L29, V30, N33, H34, E37, N38, H41, D90, I92, I93, Q96, M322, T323, P324, G325, W327, R328, D329, K352, N353, D354, R356, A385, N386, Q387, S388 and R392. The P.abr ACE2 α-helix 323-330 docks against the HKU5 RBM and forms a constellation of interactions, including R328_P.abrACE2_ forming polar interactions with the Y508_HKU5_, Y558_HKU5_ and D460_HKU5_ side chains along with pi-stacking interactions with the Y560_HKU5_ side chain and D329_P.abrACE2_ forming a salt bridge and a hydrogen bond with K544_HKU5_ and Y558_HKU5_, respectively (**Fig 2B**). Furthermore, N38_P.abrACE2_ and K352_P.abrACE2_ are hydrogen-bonded to the S547_HKU5_ side chain hydroxyl, K352_P.abrACE2_ is hydrogen-bonded to the Y545_HKU5_ and the T510_HKU5_ backbone carbonyls (**Fig 2C**). At the tip of the RBM, Y518_HKU5_ is hydrogen-bonded to R26_P.abrACE2_ and D90_P.abrACE2_ whereas K520_HKU5_ forms polar interactions with the N386_P.abrACE2_ side chain and backbone carbonyl along with the A385_P.abrACE2_ backbone carbonyl (**Fig 2C**).

Comparison of the structures of the P.abr ACE2-bound HKU5 RBD, *Pipistrellus pipistrellus* (P.pip) ACE2-bound NeoCoV RBD^39^, hDDP4-bound MERS-CoV RBD^48,49^, and hDDP4-bound HKU4^38^ RBD shows that HKU5, MERS-CoV and HKU4 utilize the same side of the RBM β-sheet to engage their cognate receptors (**Figure S3**). Conversely, NeoCoV recognizes ACE2 via the tip of the RBM, including the distal β-hairpins, which are shorter for NeoCoV and HKU5 relative to MERS-CoV and HKU4 (**Figure S3**). In spite of a conserved overall architecture of the RBMs of these four viruses, their fine molecular differences mediate the three observed binding modes: HKU5 and ACE2, NeoCoV and ACE2 as well as MERS-CoV/HKU4 and DPP4.

Analysis of amino acid residue conservation reveals a marked RBM sequence diversity among HKU5 isolates (**Fig 2D**), possibly impacting receptor engagement. To understand the impact of RBM residue polymorphism on receptor recognition, we assessed binding of the aforementioned P.abr ACE2 ectodomain construct to two distinct HKU5 RBD isolates using biolayer interferometry (BLI). We selected the HKU5-19s and HKU5-33s, which differ at positions D_HKU5-19s_460S_HKU5-33s_, Y_HKU5-19s_508H_HKU5-33s_, T_HKU5-19s_515F_HKU5-33s_, S_HKU5-19s_516T_HKU5-33s_, K_HKU5-19s_544D_HKU5-33s_, Y_HKU5-19s_545R_HKU5-33s_ and Y_HKU5-19s_558S_HKU5-33s_, due to the large number of residue substitutions. Furthermore, HKU5-33s harbors two short insertions between residues 549 and 550 (QELP) and between residues 552 and 553 (F) of the HKU5-19s RBD (**Fig 2E**). We observed that P.abr ACE2 bound with an approximately 2-fold enhanced affinity to the HKU5-19s RBD, relative to the HKU5-33s RBD, mostly driven by slower off-rates (**Fig 2F-G and Figure S4**). These findings indicate that the extensive RBM polymorphism among HKU5 isolates modulates receptor binding affinity and possibly antigenicity, as observed with SARS-CoV-2 variants^35,50–56^.

### Determinants of ACE2 species tropism

To functionally validate the role of the ACE2-interacting residues identified in our cryoEM structure, we evaluated binding of P.abr ACE2 to HKU5-19s RBD point mutants each harboring a residue substitution designed to interfere with receptor engagement. Using BLI, we observed that the HKU5-19s Y464A, Y558A, A517R and G519W substitutions abrogated binding entirely whereas the Y518A substitution did so almost completely (**Fig 3A**). Furthermore, the K520G and Y545G RBD mutations also disrupted P.abr ACE2 binding in the HKU5-19s RBD background (**Fig 3A**), possibly reducing receptor binding affinity for the HKU5 isolates in which these changes naturally occur although epistasis may modulate this effect^51^. Consistently, the HKU5-19s Y464A, A517R, Y518G, G519W, K520G, Y545G, and Y558G RBD mutants were impaired in their ability to bind P.abr ACE2 transiently expressed in HEK293T cells. Moreover, HKU5-19s S VSV pseudovirus harboring the Y464A, A517R, K520G, or Y558G substitutions exhibited the most significant reduction of entry into HEK293T cells transiently expressing P.abr ACE2 (**Fig 3B-C**).

**Figure 3.**
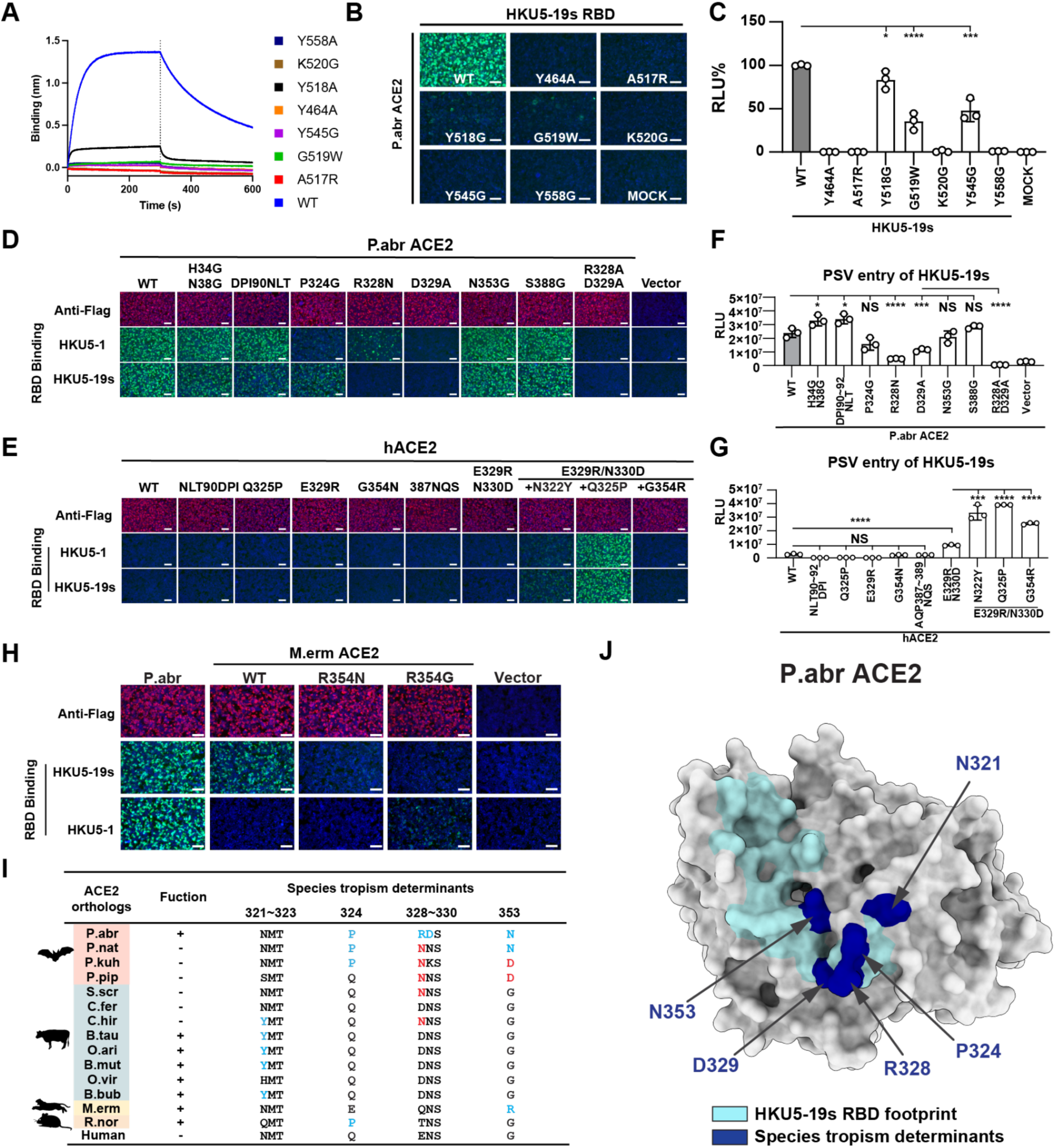
HKU5 molecular determinants of ACE2 host species tropism. **A**, Binding of the P.abr ACE2 construct comprising the peptidase and the dimerization domains (residues 20-724) to the wildtype (WT) HKU5-19s and to the listed RBD interface mutants immobilized on biolayer interferometry streptavidin (SA) biosensors. **B-C**, RBD-hFc binding (B) and pseudovirus entry (C) efficiencies of HKU5-19s mutants in HEK293T cells transiently expressing P.abr ACE2. The entry efficiency of wildtype HKU5-19s S VSV pseudovirus was set as 100%. **D-E,** HKU5-19s and HKU5-1 RBD-hFc binding to HEK293T cells transiently expressing the indicated P.abr ACE2 or hACE2 mutants assessed by immunofluorescence. **F-G,** HKU5-19s S VSV pseudovirus entry into HEK293T cells transiently expressing the indicated ACE2 mutants. **H**, HKU5-19s and HKU5-1 RBD-hFc binding to HEK293T cells transiently expressing wildtype and mutants M.erm ACE2. **I,** Summary of ACE2 residues governing species tropism for HKU5-19s. Favorable and unfavorable residues in ACE2 orthologs are highlighted in blue and red, respectively, using P.abr ACE2 residue numbering. (+): functional; (-): non-functional. **J,** Summary of the critical determinants of HKU5-19s receptor recognition. Key residues mentioned in J are highlighted in navy color on the P.abr ACE2 structure rendered as a grey surface. The rest of the HKU5-19s RBD footprint is shown in cyan. Data are shown as the MEAN ± SD of 3 biological replicates for C, F, and G. Statistical analyses used unpaired two-tailed t-tests: *: *p* < 0.05,**: *p* < 0.01, ***: *p* < 0.005, and ****: *p* < 0.001, NS: not-significant. Data representative of two independent experiments for F and G, and a single experiment for C. Scale bars: 100 μm.

Assessment of the impact of mutations of key P.abr ACE2 residues contacting the HKU5 RBD further supported these findings: the P324G, R328N (glycan knockin), D329A and R328A/D329A substitutions dampened HKU5-19s binding to and pseudovirus entry in HEK293T cells transiently transfected with these P.abr ACE2 mutants (**Fig 3D,F**). Given that several HKU5-interacting P.abr ACE2 residues are conserved or conservatively substituted in human ACE2 (hACE2, **Table 2**), we set out to determine the molecular determinants of the lack of efficient hACE2 binding and utilization (**Fig 1E-F**). Concurring with the central role of the P.abr ACE2 R328 and D329 residues, we found that a hACE2 double mutant harboring these residues (E329R/N330D) promoted detectable but weak HKU5-19s S pseudovirus entry into cells (**Fig 3E,G**). A hACE2 triple mutant harboring the E329R/N330D substitutions along with N322Y, Q325P or G354R promoted robust HKU5-19s S pseudovirus entry along with detectable binding of the RBD-Fc constructs except for the E329R/N330D/G354R (**Fig 3E,G**). Although the N321_P.abrACE2_ glycosylation sequon is conserved relative to hACE2 (equivalent to N322_hACE2_), no evidence of glycosylation could be detected in the HKU5 RBD/P.abr ACE2 complex cryoEM map (as is the case for P.abr ACE2 N38). The positioning of the N322_hACE2_ oligosaccharide in SARS-CoV-2 RBD/hACE2 structures^35,57^ suggests that it could disrupt HKU5 RBD binding, thereby explaining the improved hACE2 usage resulting from the N322Y substitution in a E329R/N330D background (**Fig 3F-G**). Unlike HKU5-19s, the HKU5-1 RBD-hFc did not bind M.erm ACE2, likely due to differences at RBD position 472: E472_HKU5-19s_ may form a salt bridge with R354_M.ermACE2_ whereas A472_HKU5-1_ could not form such contacts (**Fig 3H**). The R354N or R354G mutation significantly reduced HKU5-19s RBD-hFc binding efficiency to M.erm ACE2, highlighting the importance of the residue at position 354_hACE2/M.ermACE2_ (equivalent to 353_P.abrACE2_) for tropism determination (**Fig 3F)**.

**Table 2.**
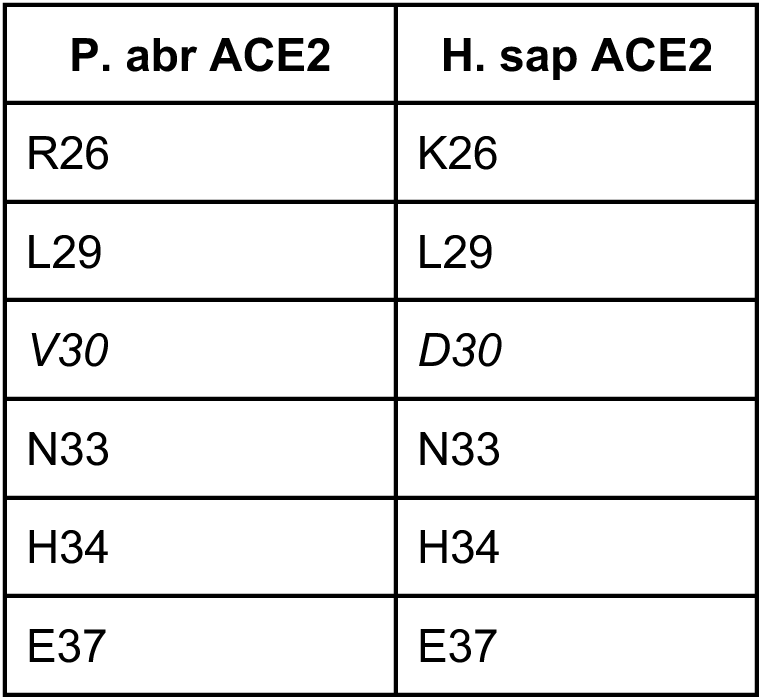
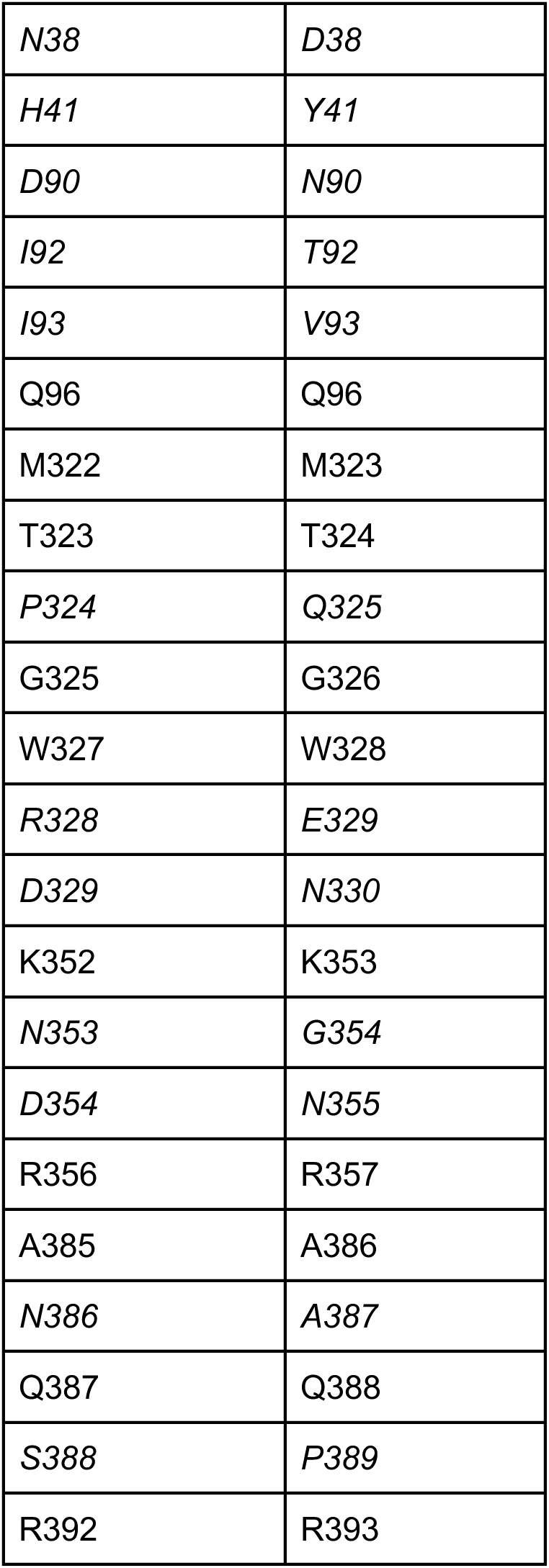
Conservation analysis of P.abr ACE2 residues recognized by HKU5 in hACE2. Non-conserved residues are italicized.

Overall, we identified and validated the functional impact of four determinants of receptor species tropism, all of which are situated in the HKU5-binding footprint, along with preferred amino acid residue for each of them (**Fig 3I-J**). These include P.abr ACE2 positions (i) 321 where a tyrosine is preferred (disrupting an N-linked glycosylation sequon), (ii) 324 where a proline is preferred, (iii) 328-329 where arginine and aspartate residues are preferred, respectively, and (iv) 353 where an asparagine or arginine is preferred by HKU5-19s.

### Identification of HKU5 countermeasures

To understand antigenic relationships among merbecoviruses, we assessed the ability of MERS-CoV infection-elicited human antibodies to inhibit HKU5-19s S VSV pseudovirus entry into HEK293T cells transiently transfected with P.abr ACE2. We used plasma collected from individuals who were hospitalized with MERS-CoV infection (prior to the COVID-19 pandemic) and selected samples at time points corresponding to peak neutralization titers against MERS-CoV EMC/2012^58^ **(Table 3)**. Neutralizing activity (reaching at least 50% inhibition of entry at 1/10 plasma dilution) was detected for three (15, 16 and 24) out of 28 plasma samples evaluated **(Fig 4A and Fig S5)**. The rare and very weak HKU5 cross-neutralization mediated by MERS-CoV-elicited polyclonal plasma antibodies underscores the genetic and antigenic distance of the S and RBD of these two viruses, sharing approximately 65% and 50% amino acid sequence identity, respectively. However, broadly neutralizing monoclonal antibodies targeting the fusion machinery (S_2_ subunit) stem helix^59,60^ (S2P6 and B6) or the fusion peptide/S_2_’ cleavage site^61,62^ (76E1) inhibited authentic HKU5-1 propagation in Caco-2/P.abr ACE2 cells **(Fig 4B)** and HKU5-19s S VSV pseudovirus **(Fig S5)**, consistent with the high degree of conservation of the targeted epitopes.

**Table 3.** Demographic information of MERS-CoV-infected individuals. DM, diabetes mellitus; HTN, hypertension; ARDS, acute respiratory distress syndrome; ESRD, end stage renal disease; IHD, ischemic heart disease; AKI, acute kidney injury; COPD, chronic obstructive pulmonary disease; BA, bronchial asthma; BPH, benign prostatic hypertrophy; Afib, atrial fibrillation.

| Patient ID | Sex | Age | Comorbidities |
| --- | --- | --- | --- |
| 1 | M | 42 | DM, HTN |
| 2 | M | 80 | ESRD, DM, HTN |
| 5 | M | 27 | DM, HTN |
| 6 | M | 64 | DM, HTN |
| 7 | M | 44 | DM |
| 8 | M | 51 | DM, HTN |
| 9 | M | 70 | DM, HTN, IHD |
| 10 | M | 35 | psychiatric illness |
| 11 | F | 55 | DM, hypothyroidism, AKI |
| 12 | M | 49 | DM |
| 13 | F | 41 | DM |
| 14 | M | 65 | DM, HTN, IHD |
| 15 | M | 46 | DM, HTN, COPD |
| 16 | M | 53 | DM, HTN |
| 17 | M | 49 | heart failure |
| 18 | M | 53 | HTN |
| 19 | M | 32 | – |
| 20 | M | 75 | DM, HTN, epilepsy, dyslipidemia |
| 21 | F | 67 | DM, HTN |
| 23 | M | 63 | DM, BA, HTN |
| 24 | M | 50 | DM |
| 26 | M | 31 | ESRD, DM, HTN |
| 27 | F | 70 | ESRD |
| 29 | M | 69 | BPH, Afib |
| 31 | F | 62 | ARDS, Alzheimer, DM, HTN |
| 32 | M | 42 | DM |
| 33 | M | 58 | – |

**Figure 4.**
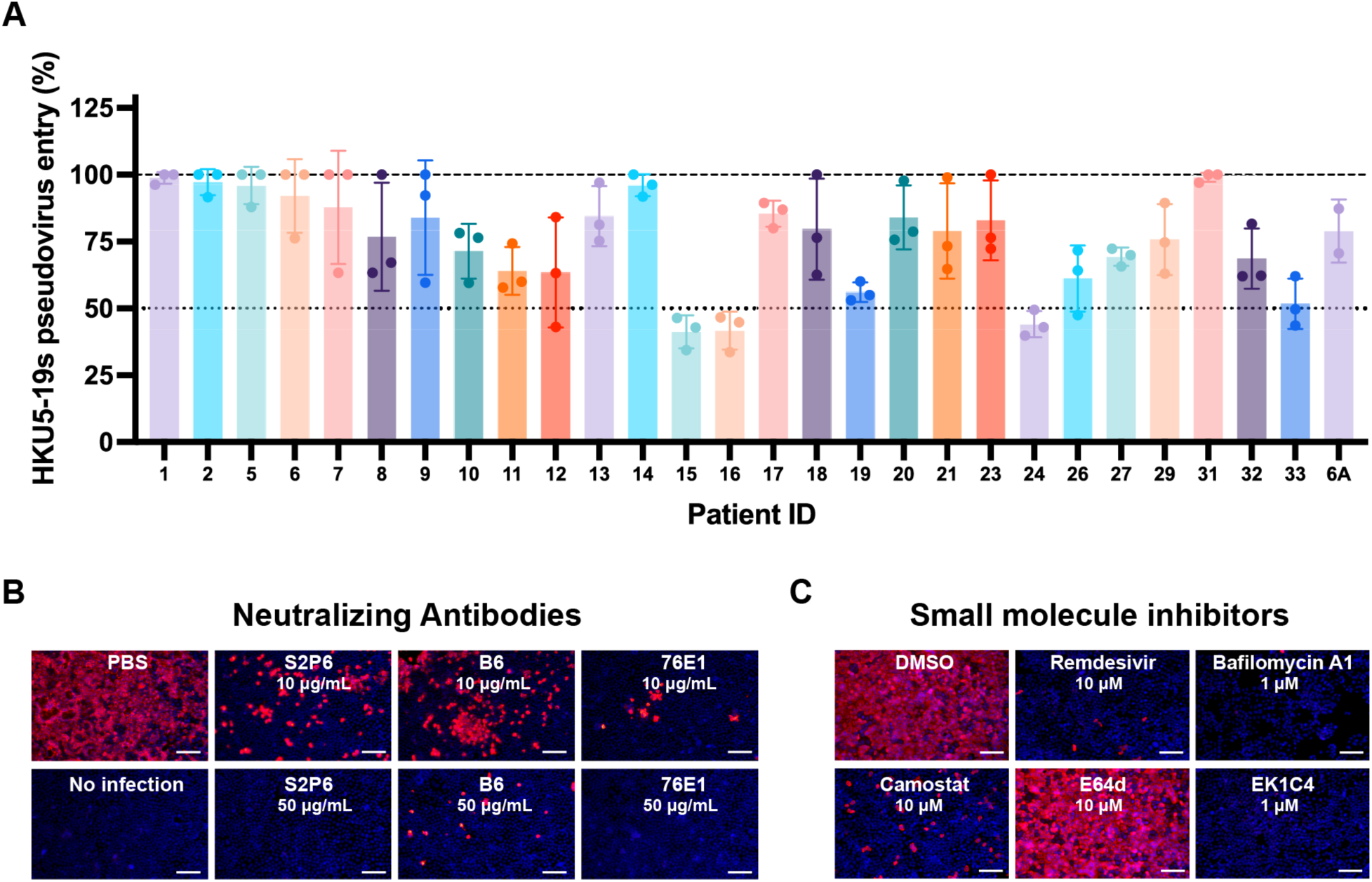
Identification of countermeasures against HKU5 merbecoviruses. **A,** Neutralization of HKU5-19s S VSV pseudovirus mediated by a panel of MERS-CoV infection-elicited human plasma^58^. Bars represent the mean of three biological replicates with SD and data points correspond to the mean of two technical replicates within each biological replicate carried out with distinct batches of pseudoviruses. **B-C**, Evaluation of inhibition of authentic HKU5-1 propagation in Caco-2 cells stably expressing P.abr ACE2 by the indicated concentration of broadly neutralizing antibodies (B) or small molecule inhibitors (C). HKU5-1 was detected by immunofluorescence using an anti-HKU5 N antibody at 24 hpi. Scale bars: 200 μm.

We subsequently screened a panel of small molecule inhibitors to evaluate their ability to block propagation of authentic HKU5-1 in Caco-2/P.abr ACE2 cells. Camostat, which inhibits TMPRSS2 and related serine proteases^63,64^, but not the E64d cysteine protease (cathepsin B/L) inhibitor, abrogated HKU5-1 propagation at a concentration of 10 µM **(Fig 4C)**. Bafilomycin A1, which hinders endosomal acidification^65^, also inhibited HKU5-1 at a concentration of 1 µM **(Fig 4C)**. These findings suggest that HKU5-1 can infect Caco-2/P.abr ACE2 cells via TMPRSS2-mediated plasma membrane fusion and via endosomal fusion, the latter process relying on non-cysteine proteases, possibly aspartyl or serine proteases. The involvement of TMPRSS2 and related TMPRSS proteases in HKU5-1 entry into Caco-2/P.abr ACE2 cells is consistent with the presence of a polybasic cleavage site at the S_1_/S_2_ junction (R_742_VRR_745_, HKU5-1 residue numbering), resulting in cleavage of the S trimers incorporated in pseudovirus during biogenesis **(Fig S6)**, which was previously described to lead to plasma membrane fusion for MERS-CoV^66,67^. Conversely, the inability of E64d to reduce HKU5-1 infection of Caco-2 cells is likely explained by the suboptimal endosomal cysteine protease site (N_760_FTS_763_), which introduces an N-linked glycosylation sequon^46,68,69^. We note that the S_1_/S_2_ junction polymorphism among HKU5 isolates impacts S processing during biogenesis (**Fig S6**), which could modulate the relative potency of each of these inhibitors. We also found that the OC43 HR2-derived EK1C4 lipopeptide fusion inhibitor^70,71^, which interferes with S fusogenic conformational changes, and remdesevir^72,73^, the inhibitor of coronavirus RNA-dependent RNA polymerase^74^, inhibited HKU5-1 propagation at a concentration of 1 µM and 10 µM, respectively **(Fig 4C)**. These findings indicate that inhibitors targeting host proteases mediating S proteolytic activation, such as TMPRSS2, or conserved coronavirus target sites, such as the S fusion machinery or viral polymerase, are promising candidates for pandemic preparedness against HKU5 and other coronaviruses.

## Discussion

Phylogenetic classification of merbecovirus RBDs underscores the exceptional diversity of these pathogens harboring RBDs sharing as little as 30% amino acid sequence identity which cluster in at least six clades. The identification of DPP4 as the host receptor for members of the MERS-CoV clade upon its emergence in 2012^36^ and subsequently for members of the HKU4 clade^37,38,77^, led to the assumption that DPP4 is a universal receptor for merbecoviruses. The discovery of bat-borne merbecoviruses in Africa utilizing a broad spectrum of ACE2 orthologs^39,40^ indicated that receptor diversity was greater than previously appreciated for these viruses. The discovery that viruses from the HKU5 clade and the MOW15-22 clade^44^ use the ACE2 receptor profoundly alters our understanding of merbecovirus evolution and shows that ACE2 usage is a broadly shared property among several clades. Furthermore, the structural specialization of the RBM within each clade along with the entirely distinct ACE2 binding modes lead to recognition of receptor regions located up to 50Å apart. These findings indicate that ACE2 utilization has been independently acquired at least three times among merbecoviruses (NeoCoV/PDF-2180, MOW15-22/PnNL2018B, HKU5) and at least twice among other coronaviruses (SARS-CoV-1/SARS-CoV-2^35,57,78^ and NL63^75^, **Fig 5**). We note that utilization of amino-peptidase N among a large set of coronaviruses also results from at least three convergent evolution events for TGEV/CCoV-HuPn-2018^79,80^, PDCoV^81^ and 229E^82^. Recurrent coronavirus convergence to use a small number of host transmembrane protease receptors, for which the proteolytic activity does not participate in entry, might result from specific size and geometrical requirements for infection or of the clustering of receptors in specialized membrane patches^45^.

**Figure 5.**
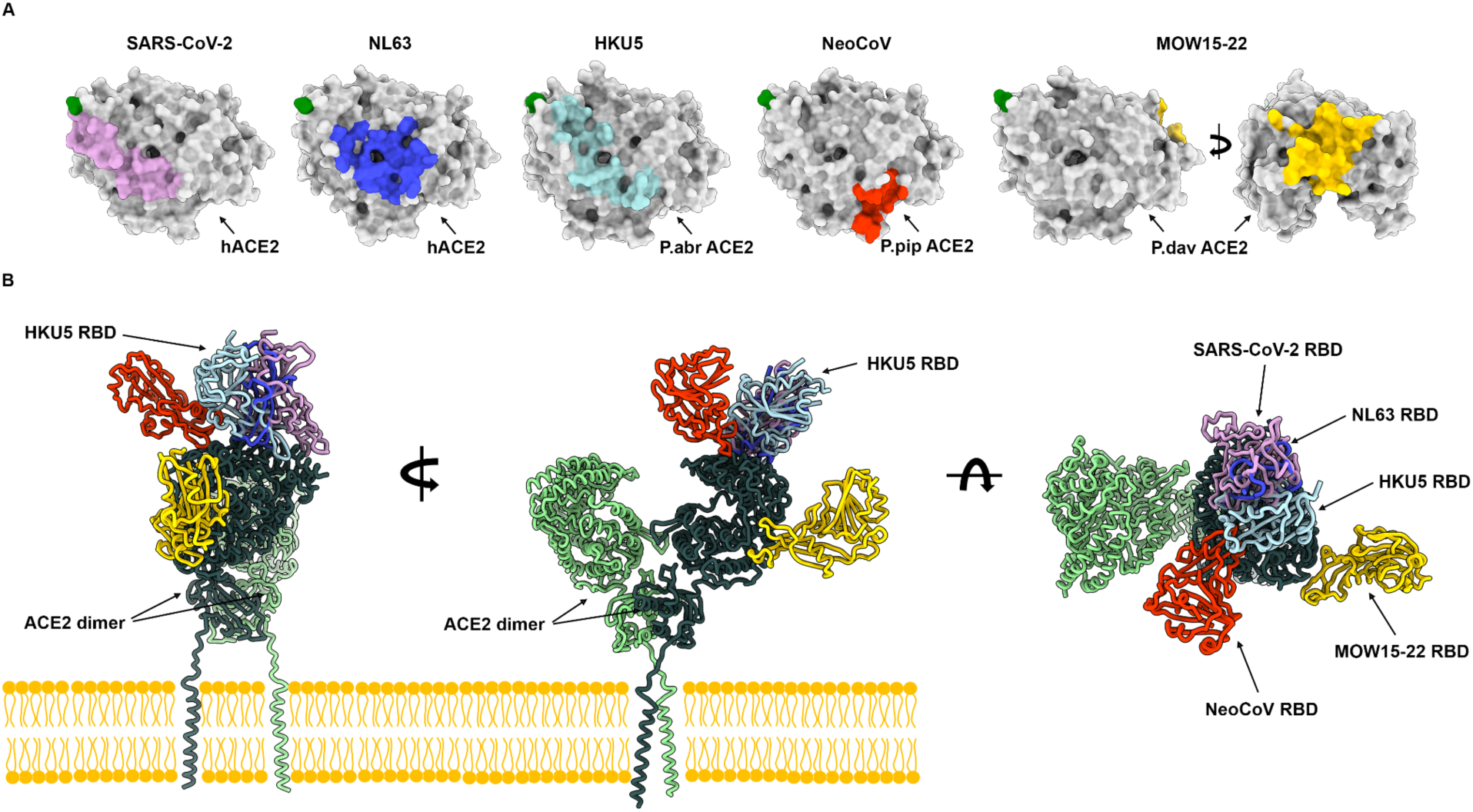
Coronaviruses have evolved ACE2 utilization at least five times independently. **(A)** RBD footprints of ACE2-using coronaviruses on their cognate receptors. **(B)** Comparison of the binding modes of the SARS-CoV-2 (PDB 7TN0)^35^, NL63 (PDB 3KBH)^75^, HKU5 RBDs, NeoCoV (PDB 7WPO)^39^ and MOW15-22 (PDB 9C6O)^44^ to bat ACE2 (not shown for clarity) or hACE2 (PDB 6M1D, B0AT1 not shown for clarity)^76^.

Given that the RBD is the main target of coronavirus neutralizing antibodies^26,30,58,83^, the rare and weak cross-neutralization of HKU5 observed with MERS-CoV infection-elicited polyclonal plasma antibodies concurs with the extensive genetic divergence among merbecovirus RBD clades. These findings are reminiscent of the weak cross-clade neutralization observed for sarbecoviruses^84,85^ which harbor RBDs much more similar to each other than merbecovirus RBDs. As a result, the development of vaccines with pan-merbecovirus neutralizing activity may prove to be exceptionally challenging^58,86^. However, our data suggest that rare monoclonal antibodies with broadly neutralizing merbecovirus activity were present in the plasma of a few subjects in the cohort analyzed, explaining the detectable (albeit weak) cross-neutralization and motivating future isolation and characterization of these antibodies. We hypothesize that these rare antibodies may target conserved core RBD antigenic sites or conserved fusion machinery epitopes^59,60,62,87–90^.

Although we could not detect hACE2-mediated HKU5 RBD binding or S VSV pseudovirus entry, exogenous trypsin addition enabled weak propagation of authentic HKU5-1 in Caco-2 human cells, in line with a previous study^46^. Given the extensive RBM polymorphism observed in the HKU5 clade, possibly resulting from immune pressure in the reservoir host, and our limited understanding of the genetic diversity of these viruses (due to sparse sampling of merbecoviruses in bats), HKU5 may acquire more efficient hACE2 usage and constitute a zoonotic threat due to the lack of merbecovirus immunity in the human population. We provide a blueprint for monitoring HKU5 residue substitutions at the interface with the molecular determinants of ACE2 species tropism delineated here to detect possible future HKU5 adaptations to the human receptor. The identification of several inhibitors of HKU5-1 propagation in human cells, including camostat (a clinical inhibitor of host serine proteases) and remdesevir (an FDA-approved inhibitor of coronavirus RNA-dependent RNA polymerase), suggests that these compounds could be used in case of possible future outbreaks.

## Author contributions

YJP, CL, HY and DV conceived the project. YJP and MAT designed glycoprotein constructs. JTB, and CS recombinantly expressed glycoproteins. CL and CBM cloned RBD-hFc and ACE2 mutants and conducted RBD-hFc binding assays. JTB. conducted biolayer interferometry binding experiments. CL, JL, and CBM carried out pseudovirus entry and neutralization assays. YJP carried out cryoEM sample preparation, data collection, and processing. YJP and DV built and refined the structure. PL performed authentic virus infection and inhibition assays. HY and DV wrote the manuscript with input from all authors. YJP, CL, JL, AAd, HY and DV analyzed the data. AAl contributed unique reagents. CJC, TNS and CL conducted phylogenetic analysis.

## Acknowledgments

This study was supported by the National Institute of Allergy and Infectious Diseases (P01AI167966 to TNS and DV, DP1AI158186 and 75N93022C00036 to D.V.), the National Institute of General Medical Sciences (T32GM141848 to C.J.C.), an Investigators in the Pathogenesis of Infectious Disease Awards from the Burroughs Wellcome Fund (D.V.), a Searle Scholars Award (T.N.S.) the University of Washington Arnold and Mabel Beckman cryoEM center and the National Institute of Health grant S10OD032290 (to DV). This study was also supported by the National Natural Science Foundation of China (NSFC) projects (82322041, 32270164, 32070160 to H.Y., 323B2006 to C.B.M.), the National Key R&D Program of China (2023YFC2605500 to H.Y.), the Fundamental Research Funds for the Central Universities (2042023kf0191, 2042022kf1188 to H.Y.), Natural Science Foundation of Hubei Province (2023AFA015 to H.Y.). DV is an Investigator of the Howard Hughes Medical Institute and the Hans Neurath Endowed Chair in Biochemistry at the University of Washington. We thank the University of Utah Center for High Performance Computing supported by NIH (1S10OD021644-01A1). We thank Lu Lu (Fudan University) for providing EK1C4 peptides. We thank Qiang Ding (Tsinghua University), Zheng-Li Shi (Wuhan Institute of Virology), and Qihui Wang (CAS Key Laboratory of Pathogenic Microbiology & Immunology, China) for sharing some mammalian ACE2-expressing plasmids.

## Data and code availability

The cryoEM maps and model have been deposited to the electron microscopy data bank and protein data bank with accession numbers EMD-46512, PDB-9D32. The pipeline for phylogenetic analysis is available from GitHub: https://github.com/tstarrlab/MERSr_phylo.

## Methods

### Cells

Cell lines used in this study were DH10B competent cells (Thermo Fisher Scientific), HEK293T (ATCC, CRL-11268), and Caco-2 (HTB-37). HEK293T and Caco-2 cells were cultured in 10% FBS (Fisher Scientific-Cytiva), 1% penicillin-streptomycin (Thermo Fisher Scientific) DMEM at 37℃, 5% CO_2_. I1-Hybridoma cell line producing a neutralizing antibody targeting the VSV glycoprotein (VSV-G) was cultured in Minimum Essential Medium (MEM) with Earles’s balanced salts, 2.0 mM of L-glutamine (Gibico), and 10% FBS. HEK293T or Caco-2 stable cell lines overexpressing various receptors were generated using lentivirus transduction and selected and maintained in the growth medium with puromycin (1 μg /ml).

### Construct design

The wildtype P.abr (Pipistrellus abramus) ACE2 ectodomain encoding residues 18-724 (ACT66266.1) was inserted after the N-terminal signal peptide MPMGSLQPLATLYLLGMLVASVLA and C-terminally fused to a thrombin cleavage site followed by a SSGGS linker, an avi tag, a GGS short linker and an octa-histidine tag for affinity purification.. The HKU5-19s wildtype (AGP04932.1) spike RBD encoding residues 390-587 and the Y558A, K520G, Y518A, Y464A, Y545G, G519W, A517R receptor-interface mutants, contain the N-terminal signal peptide MGILPSPGMPALLSLVSLLSVLLMGCVAETGT, followed by a thrombin cleavage site, a SSGGS flexible linker, an avi tag, a GGS flexible linker and a C-terminal octa-histidine tag. HKU5-33s wildtype (AGP04943.1) spike RBD encoding residues 390-592 contain the N-terminal signal peptide MGILPSPGMPALLSLVSLLSVLLMGCVAETGT, followed by a thrombin cleavage site, an avi tag, a short GGS flexible linker and a C-terminal octa-histidine tag. All the above genes were codon-optimized for expression in mammalian cells, synthesized, and inserted in the pcDNA3.1(+) by Genscript. Plasmids expressing wild-type (WT) or mutated bat and non-bat mammalian ACE2 orthologs were constructed by inserting human codon-optimized sequences with/without specific mutations into a lentiviral transfer vector (pLVX-EF1a-Puro, Genewiz) with C-terminus 3×FLAG tags (DYKDHD-G-DYKDHD-I-DYKDDDDK) and single FLAG tag (DYKDDDDK) for non-bat mammalian ACE2 orthologs. For pseudovirus production, human codon-optimized spike sequences of HKU5-19s (AGP04932.1), HKU5-1 (YP_001039962) fused C-terminal HA tag (YPYDVPDYA) were cloned into the pCAGGS vector with C-terminal deletions (residues 13-15) for improving the pseudovirus assembly efficiency^91^. The DNA fragments for cloning ACE2 chimera or RBD mutants were generated by overlap extension PCR or gene synthesis and verified by commercial DNA sequencing. For the expression of recombinant CoVs RBD-hFc fusion proteins, plasmids were constructed by inserting HKU5-19s RBD (residues 385-586), HKU5-1 RBD (residues 385-586), coding sequences into the pCAGGS vector containing an N-terminal CD5 secretion signal peptide (MPMGSLQPLATLYLLGMLVASVL) and C-terminal hFc-twin-strep-3xFLAG tag (WSHPQFEKGGGSGGGSGGSAWSHPQFEKGGGRSDYKDHDGDYKDHDIDYKDDDDK) for purification and detection. Heavy chain and light chain genes encoding 76E1 IgG were inserted into pcDNA3.1(+) plasmid.

### Recombinant protein production

The P. abramus ACE2 ectodomain, the wildtype HKU5-19s and HKU5-33s RBDs along with HKU5-19s interface point mutants were expressed in Expi293F cells (Thermo) maintained at 37°C and 8% CO_2_. Cells were transfected using Expifectamine293 (Thermo) following the manufacturer’s protocol. Four to five days post-transfection, Expi293 cell supernatant was clarified by centrifugation at 4,121g for 30 minutes. Supernatants were supplemented to a final concentration of 100 mM Tris pH 8.0, and 300 mM NaCl before binding to Ni excel resin (Cytiva) previously equilibrated in 100 mM Tris pH 8.0 or 100 mM Tris pH 7.4 and 300 mM NaCl. Nickel resin was washed with 100mM Tris pH 8.0 or 100mM Tris pH 7.4, 300 mM NaCl, and 10mM imidazole prior to elution with 100mM Tris pH 8.0 or 100mM Tris pH 7.4, 300 mM NaCl, and 300mM imidazole.

ACE2 ectodomains were concentrated using centrifugal filter devices with a MWCO of 30kDa and eluted on a Superdex200 increase 10/300 size-exclusion column (Cytiva) equilibrated in 50mM Tris pH 7.4 and 150 mM NaCl and monodisperse fractions were pooled and used immediately or flash frozen and stored at -80℃ until use. For structural characterization, purified HKU5-19s RBD was concentrated using a centrifugal filter device with a MWCO of 10kDa and eluted on a Superdex200 increase 10/300 size-exclusion column (Cytiva) equilibrated in 50mM Tris pH 7.4 and 150 mM NaCl and fractions containing monomeric RBD were used immediate or flash frozen and stored at -80C until use.

For BLI, purified RBDs were biotinylated using the BirA biotin-protein ligase reaction kit (Avidity) after buffer exchanging into 50 mM Tris pH 7.4 and 150 mM NaCl using a centrifugal filter device with a MWCO of 10kDa. Affinity purification as above-described was carried out a second time to get rid of BirA and free biotin. Purified biotinylated HKU5-19s interface point mutant RBDs were buffer exchanged into 50 mM Tris pH 7.4 and 150 mM NaCl and used immediately or flash frozen. Purified biotinylated HKU5-19s and HKU5-33s RBD were concentrated using a centrifugal filter device with a MWCO of 10kDa and run on a Superdex200 increase 10/300 size-exclusion column (Cytiva) equilibrated in 50mM Tris pH 7.4 and 150 mM NaCl and fractions containing monomeric RBD were flash frozen and stored at -80℃ until use.

HEK293T cells were transfected with HKU5-19s WT or mutant RBD plasmids using GeneTwin reagent (Biomed, TG101-01). Subsequently, the culture medium of the transfected cells was replenished with the SMM 293-TII Expression Medium (Sino Biological, M293TII) 4-6 hours post-transfection, and the protein-containing supernatant was collected every two days for 2-3 batches. Antibodies and recombinant RBD-hFc proteins were purified using Pierce Protein A/G Plus Agarose (Thermo Scientific, 20424). In general, Fc-containing proteins were enriched by the agarose, washed with wash buffer (100 mM Tris/HCl, pH 8.0, 150 mM NaCl, 1 mM EDTA), eluted using the Glycine buffer (100 mM in H_2_O, pH 3.0), and immediately neutralized with 1/10 volume of 1M Tris-HCI, pH 8.0 (15568025, Thermo Scientific). Proteins with twin-strep tag were purified using Strep-Tactin XT 4Flow high-capacity resin (IBA, 2-5030-002), washed by wash buffer (100 mM Tris/HCl, pH 8.0, 150 mM NaCl, 1 mM EDTA), and then eluted with buffer BXT (100 mM Tris/HCl, pH 8.0, 150 mM NaCl, 1 mM EDTA, 50 mM biotin). All eluted proteins were concentrated using Ultrafiltration tubes, buffer-changed to PBS, and stored at -80℃. Protein concentrations were determined by the Omni-Easy Instant BCA Protein Assay Kit (Epizyme, ZJ102) or BLI assays using Octet RED96 instrument (Molecular Devices).

To express the 76E1 monoclonal antibody, Expi293F cells were transiently transfected with plasmids encoding for the IgG heavy and light chains (with an equal ratio of each plasmid) using Expifectamine following the manufacturer’s protocol. Four days after transfection, Expi293F cell supernatant was clarified by centrifugation at 4,121xg for 30 minutes and diluted with 20mM phosphate pH 8.0 and 0.5 mM PMSF. Supernatant was passed through a Protein A affinity column (Cytiva) twice before being washed with 20 mM phosphate pH 8.0. 76E1 was eluted with 100 mM citric acid pH 3.0 into tubes containing 1M Tris pH 9.0 for immediate neutralization of the low pH used for elution. Eluted 76E1 was buffer-exchanged into 20 mM phosphate buffer pH 8.0 and 100 mM NaCl and the purity was confirmed by SDS-PAGE.

To express the B6 monoclonal antibody, ExpiCHO cells were transiently transfected with plasmids encoding for the IgG heavy and light chains (with an equal ratio of each plasmid) following the manufacturer’s protocol using Expifectamine CHO. Eight days after transfection, ExpiCHO cell supernatant was clarified by centrifugation at 3,500xg for 30 minutes then filtered via vacuum through a 0.2 µm aPES membrane, supplemented with a buffer containing 500 mM sodium phosphate pH 8.0 to a final concentration of 20 mM and PMSF added to a final concentration of 0.1 mM. Supernatant was passed over a Protein A affinity column (Cytiva) twice before being washed with 20 mM sodium phosphate pH 8.0. B6 was eluted with 100 mM citric acid pH 3.0 into tubes containing 1M Tris pH 9.0 for immediate neutralization to a final pH of 8.0. The eluted B6 antibody was buffer-exchanged into 25 mM phosphate pH 8.0, 100 mM NaCl and purity was confirmed by SDS-PAGE in both reducing and non-reducing conditions.

### RBD-hFc live-cell binding assay

RBD-hFc live-cell binding assays were conducted following a previously described protocol. The coronavirus RBD -hFc recombinant proteins were diluted in DMEM at 4 μg/mL and incubated with HEK293T cells transiently expressing different ACE2 for 30 minutes at 37℃ at 36 hours post-transfection. Subsequently, cells were washed once with Hanks’ Balanced Salt Solution (HBSS) and incubated with 1 μg/mL of Alexa Fluor 488-conjugated goat anti-human IgG (Thermo Fisher Scientific; A11013) and Hoechst 33342 (1:10,000 dilution in HBSS) diluted in HBSS/1% BSA for 1 hour at 37℃. After another round of washing with HBSS, the cells were incubated with fresh HBSS. The images were captured using a fluorescence microscope (MI52-N). The relative fluorescence intensities (RFUs) of the stained cells were determined by a Varioskan LUX Multi-well Luminometer (Thermo Scientific).

### Pseudovirus production and entry assays for receptor identification

VSV-dG-based pseudovirus (PSV) carrying trans-complementary S glycoproteins from various coronaviruses were produced following a modified protocol as previously described. Briefly, HEK293T cells were transfected with coronaviruses S glycoproteins expression plasmids. At 24 hours post-transfection, cells were infected with 1.5×10^6^ TCID50 VSV-G glycoprotein-deficient VSV expressing GFP and firefly luciferase (VSV-dG-fLuc-GFP, constructed and produced in-house) diluted in DMEM with 8 μg/mL polybrene and for 4-6 hours at 37 ℃. After three PBS washes, the culture medium was replenished with fresh DMEM or SMM 293-TII Expression Medium (Sino Biological, M293TII), along with the presence of the neutralizing antibody (from I1-mouse hybridoma) targeting the VSV-G to eliminate the background produced by the remaining VSV-dG-fLuc-GFP. Twenty-four hours later, the pseudovirus containing supernatant was clarified through centrifugation at 12,000 rpm for 5 minutes at 4℃, aliquoted, and stored at -80℃. The TCID_50_ of the PSV was calculated using the Reed-Muench method^92^.

HEK293T or Caco-2 cells transiently or stably expressing the indicated ACE2 orthologues were infected by the single-round pseudovirus. Approximately 3×10^4^ trypsinized cells were incubated with pseudovirus (2×10^5^ TCID50/100 μL) in a 96-well plate to facilitate attachment and viral entry simultaneously. Before inoculation, pseudoviruses were typically treated with 100 μg/mL TPCK-trypsin (Sigma-Aldrich, T8802). Specifically, pseudoviruses produced in serum-free SMM 293-TII Expression Medium were incubated with TPCK-treated trypsin for 10 minutes at room temperature, and the proteolytic activity was neutralized by FBS in the culture medium. I1 Intracellular luciferase activity (Relative light units, RLU) was measured using the Bright-Glo Luciferase Assay Kit (Promega, E2620) and detected with a GloMax 20/20 Luminometer (Promega) or Varioskan LUX Multi-well Luminometer (Thermo Fisher Scientific) at 18 hours post-infection.

### Pseudovirus production and entry for neutralization assays

HKU5-19s S VSV pseudoviruses were produced using HEK293T cells seeded on BioCoat Cell Culture Dish: poly-D-Lysine 100 mm (Corning). Cells were transfected with respective S constructs using Lipofectamine 2000 (Life Technologies) in Opti-MEM transfection medium. After 5h of incubation at 37 °C with 5% CO_2_, cells were supplemented with DMEM containing 10% of FBS. On the next day, cells were infected with VSV (G*ΔG-luciferase) for 2h, followed by five time wash with DMEM medium before addition of anti-VSV G antibody (I1-mouse hybridoma supernatant diluted 1:40, ATCC CRL-2700) and medium. After 18-24 h of incubation at 37 °C with 5% CO_2_, pseudoviruses were collected and cell debris was removed by centrifugation at 3,000g for 10 min. Pseudoviruses were further filtered using a 0.45 µm syringe filter and concentrated 10x prior to storage at -80°C.

For plasma and monoclonal antibody neutralization assay, HEK293T cells were transfected with full-length P. abramus ACE2 using Lipofectamine 2000 (Life Technologies) in Opti-MEM transfection medium. After 5h of incubation at 37 °C with 5% CO_2_, cells were seeded into 96-well plates precoated with poly-L-lysine solution (Sigma, P4707). The following day, a half-area 96-well plate (Greiner) was prepared with 3-fold serial dilutions of each plasma (starting dilutions of 1:10, 22µL in total per well) or of the B6 and 76E1 monoclonal antibodies (starting dilutions of 500 µg/mL). An equal volume of DMEM with diluted pseudoviruses was added to each well. The pseudovirus was diluted to 1:15 to reach a target entry of ∼10^6^ relative luciferase units (RLUs). The mixture was incubated at room temperature for 45-60 minutes. 40 μL from each well of the half-area 96-well plate containing sera and pseudovirus were transferred to the 96-well plate seeded with cells and incubated for 1h. After 1h, an additional 40 μL of DMEM supplemented with 20% FBS and 2% PenStrep was added to the cells. After 18–20h of incubation at 37 °C with 5% CO_2_, 40 μL of One-Glo-EX substrate (Promega) was added to each well and incubated on a plate shaker in the dark for 5 min before reading the luciferase signal using a BioTek Neo2 plate reader. RLUs were plotted and normalized in Prism (GraphPad): 100% neutralization being cells in the absence of pseudovirus and 0% neutralization being pseudovirus entry into cells without plasma. Prism (GraphPad) nonlinear regression with “log[inhibitor] versus normalized response with a variable slope” was used to fit the curve. The mean percent pseudovirus entries at 1:10 sera dilution point are shown from the two biological replicates per sample-pseudovirus pair. Demographics of the sera samples were previously described^58^.

### Immunofluorescence assay

The expression levels of ACE2 orthologs with C-terminal fused FLAG tags were determined through immunofluorescence assays. Specifically, the transfected cells were fixed and permeabilized by incubation with 100% methanol for 10 minutes at room temperature. Subsequently, the cells were incubated with a mouse antibody M2 (Sigma-Aldrich, F1804) diluted in PBS/1% BSA for one hour at 37℃. After once PBS wash, the cells were incubated with Alexa Fluor 594-conjugated goat anti-mouse IgG (Thermo Fisher Scientific, A32742) secondary antibody diluted in 1% BSA/PBS for one hour at 37℃. The images were captured and merged with a fluorescence microscope (Mshot, MI52-N) after the nucleus was stained blue with Hoechst 33342 reagent (1:5,000 dilution in PBS).

### Western blot

Western blot assays were conducted to examine the S glycoprotein proteolytic processing and incorporation efficiency, the PSV-containing supernatant was concentrated using a 30% sucrose cushion (30% sucrose, 15 mM Tris-HCl, 100 mM NaCl, 0.5 mM EDTA) at 20,000×g for 1 hour at 4℃. The concentrated virus pellet was resuspended in 1×SDS loading buffer and incubated at 95℃ for 30 minutes, followed by western blot detecting the S glycoproteins by C-terminal HA tags and with the VSV-M serving as a loading control. The blots were then washed three times by PBST and then visualized using an Omni-ECL Femto Light Chemiluminescence Kit (EpiZyme, SQ201) by a ChemiDoc MP Imaging System (Bio-Rad).

### Cryo-electron microscopy data collection, processing and model building

The P.abr ACE2 ectodomain-bound HKU5-19s (non-biotinylated) RBD complex was prepared by mixing at 1:1.2 molar ratio followed by a 1 hour incubation at room temperature. Specimen vitrification followed two methods. 3µL of 4mg/ml complex with 6 mM 3-[(3-Cholamidopropyl)dimethylammonio]-2-hydroxy-1-propanesulfonate (CHAPSO) were applied onto freshly glow discharged R 2/2 UltrAuFoil grids^93^ prior to plunge freezing using a vitrobot MarkIV (ThermoFisher Scientific) with a blot force of 0 and 5.5 sec blot time at 100% humidity and 22°C. 3µL of 0.2 mg/mL complex without detergent was added to the glow discharged side of R 2/2 UltrAuFoil grids and 1µL was added to the back side before plunging into liquid ethane using a GP2 (Leica) with 6 sec blot time. The data were acquired using an FEI Titan Krios transmission electron microscope operated at 300 kV and equipped with a Gatan K3 direct detector and Gatan Quantum GIF energy filter, operated in zero-loss mode with a slit width of 20 eV. Automated data collection was carried out using Leginon^94^ at a nominal magnification of 105,000× with a pixel size of 0.843 Å. The dose rate was adjusted to 9 counts/pixel/s, and each movie was acquired in counting mode fractionated in 100 frames of 40 ms. A total 11,223 micrographs were collected with a defocus range between -0.2 and -3 μm and stage tilt angle of 0°, 20°, 25° and 30°. Movie frame alignment, estimation of the microscope contrast-transfer function parameters, particle picking, and extraction were carried out using Warp^95^. Particles were extracted with a box size of 120 pixels with a pixel size of 1.686Å. Two rounds of reference-free 2D classification were performed using cryoSPARC^96^ to select well-defined particle images. Initial model generation was carried out using ab-initio reconstruction in cryoSPARC and the resulting maps were used as references for heterogeneous 3D refinement. Particles belonging to classes with the best resolved HKU5 RBD and ACE2 density were selected. To further improve the data, the Topaz model^97^ was trained on Warp-picked particle sets belonging to the best classes after 2D classification and particles picked using Topaz were extracted and subjected to 2D-classification and heterogenous 3D refinements. The two different particle sets from the Warp and Topaz picking strategies were merged and duplicates were removed using a minimum distance cutoff of 95Å. After two rounds of ab-initio reconstruction-heterogeneous refinements, 3D refinement was carried out using non-uniform refinement in cryoSPARC^98^. The dataset was transferred from cryoSPARC to Relion^99^ using the pyem program package^100^ and particle images were subjected to the Bayesian polishing procedure implemented in Relion^101^ during which particles were re-extracted with a box size of 320 pixels and a pixel size of 1.0 Å. To further improve the map quality, ab-initio reconstruction in cryoSPARC was used to classify the data in three bins and the generated models were used as references for heterogeneous 3D refinement. The final 3D refinements were carried out using non-uniform refinement along with per-particle defocus refinement in cryoSPARC to yield the final reconstruction at 3.1 Å resolution comprising 550,684 particles. Reported resolutions are based on the gold-standard Fourier shell correlation (FSC) of 0.143 criterion and Fourier shell correlation curves were corrected for the effects of soft masking by high-resolution noise substitution^102,103^. Local resolution estimation, filtering, and sharpening were carried out using cryoSPARC. UCSF Chimera^104^, Coot^105^, AlphaFold3^106^ and Rosetta^107,108^ were used to fit, build, refine and relax the model into the sharpened and unsharpened cryoEM maps before validation using Phenix^109^, Molprobity^110^, EMRinger^111^ and Privateer^112^.

### Biolayer interferometry

P. abramus ACE2 binding kinetics to the HKU5-19s and HKU5-33s RBDs was assessed by BLI using an Octet RED96 instrument (Sartorius) and the Octet Data acquisition software. All BLI measurements were performed at 30°C and shaking at 1,000 rpm. Biotinylated HKU5 RBDs were diluted to 10µg/mL in 10x Octet kinetics buffer (Sartorius) and loaded onto hydrated streptavidin (SA) biosensors to 1 nm shift, equilibrated in 10x Octet kinetics buffer for 60 seconds, and dipped into P. abramus ACE2 at 300nM, 100nM, 33.3nM and 11.1nM for 900s to observe association. Dissociation was observed by dipping biosensors in a 10x Octet kinetics buffer for 300s. Baseline subtraction was done by subtracting the response from unloaded SA tips dipped into 300nM P. abramus ACE2. In addition, association phases were aligned to 0 seconds and 0 response in Octet Data Analysis HT software. Apparent dissociation constants were determined by plotting the concentration of ACE2 versus the average responses from 890-895s and fitting a one site–specific binding nonlinear fit in GraphPad Prism10. Sensorgrams were plotted in GraphPad Prism10.

P. abramus ACE2 binding to wildtype and interface mutant HKU5-19s RBDs was assessed by by loading each RBD onto hydrated streptavidin biosensors to a 1nm shift, equilibrated in 10x Octet kinetics buffer for 60 seconds, before dipping into P. abramus ACE2 at 100nM for 300s to observe association. Dissociation was observed by dipping biosensors in 10x Octet kinetics buffer for 300s. Baseline subtraction was done by subtracting the response from unloaded SA tips dipped into 100nM P. abramus ACE2. In addition, association phases were aligned to 0 seconds and 0 response in Octet Data Analysis HT software. Sensorgrams were plotted in GraphPad Prism10.

#### Authentic HKU5 infection assay

For HKU5-1(isolate LMH03F) infection-related experiments, Caco-2 cells with or without human or P.abr ACE2 stable expression were initially seeded in 96-well plates and washed with DMEM before inoculation, either in the presence or absence of indicated concentration of trypsin. Following a one-hour incubation of the indicated MOI of HKU5-1 at 37℃, the cells were washed with DMEM and further incubated for the indicated hours at 37℃. For antibodies and EK1C4 neutralization assay, HKU5-1 was preincubated with the antibodies or peptides for 1 hour at 37℃ before adding to the cells. For other small molecule inhibitors, HKU5-1 was mixed with the inhibitors and immediately incubated with the cells. After 1-hour inoculation, cells were further incubated with culture medium containing indicated concentrations of inhibitors. For immunofluorescence assays, cells were washed once by DMEM and then fixed with 4% paraformaldehyde (PFA) for 20 minutes at room temperature, permeabilized by 0.1% Triton X-100 at 25℃ for 15 minutes, and blocked by 1% BSA at 37℃ for 30 minutes at indicated time points. The expression of HKU5 N proteins was detected by rabbit anti-HKU5 N protein serum (diluted at 1:4000), followed by DL594-conjugated goat anti-rabbit IgG (Thermo, 1:1000) staining. HKU5-1 titers were determined in Caco-2/P.abr ACE2 cells infected with serially-diluted inocula and calculated using the Reed-Muench method.

### Merbecovirus RBD and mammalian ACE2 phylogenetic analysis

A dataset of merbecovirus S glycoproteins was gathered based on literature search^113–117^ followed by additional BLAST searches of NCBI databases. Identical S amino acid sequences were pared via CD-HIT^118^ and aligned via the structure-guided approach of MAFFT-DASH^119^, which utilized the PDB codes 3JCL, 5I08, 5KWB, 5X59, 6B3O, 6OHW, 6PZ8, 6Q04, and 6VSJ during alignment. RBD sequences (MERS-CoV EMC-2012 reference numbering Q377-A591, Genbank Accession JX869059) were parsed from the alignment and further pared via CD-HIT to eliminate redundant sequences with 100% RBD amino acid sequence identity or to remove sequences with ambiguous base calls, leaving 71 unique RBD sequences. The full set of sequences, along with accession numbers, are available from GitHub: https://github.com/tstarrlab/MERSr_phylo/blob/main/unaligned_sequences/GenBankAccessions.xlsx. The RBD sequence alignment was determined to be free of detectable recombination via the likelihood-based model selection program GARD^120^. The final RBD sequence set was aligned again via MAFFT-DASH (which utilized PDBs 4KQZ, 4L3N, 4QZV, 4XAK, 5XGR, 6C6Z, and 6PZ8) and combined with MHV (ACN89763), OC43 (KX344031) and HKU1 (KF686346) RBD sequences as embecovirus outgroups, with final RBD alignment available from GitHub: https://github.com/tstarrlab/MERSr_phylo/blob/main/merbeco_alignments/merbeco_RBD_aligne <u>d_v2.fasta</u>. The maximum likelihood phylogenetic relationship among RBD sequences was inferred using RAxML^121^ with the WAG+G evolutionary model, and visualized based on rooting on the embecovirus outgroups. The maximum likelihood tree originally placed a poorly supported paraphyletic relationship for the HKU25-related RBD clade (**Fig 1A**). This paraphyletic relationship is unstable (i.e., has fluctuated as we iterated on the phylogeny when adding newly described taxa); disagrees with the most parsimonious placement of an indel (deletion of equivalent MERS-CoV residues 543-552 that is specific to the HKU25-related RBDs); and disagrees with monophyletic HKU25-related clade grouping from phylogenies based on whole genome and full S sequences^122,123^ which are more confident due to longer alignment length. For our final phylogeny, we therefore enforced a constraint so that RBDs from HKU25-related viruses form a monophyletic clade. The full pipeline and intermediate analysis files are available from GitHub: https://github.com/tstarrlab/MERSr_phylo. Sequence alignments of different ACE2 orthologs or HKU5 S_1_/S_2_ regions were performed with the MUSCLE^124^ algorithm in MEGA-X^125^ (version 10.1.8). The logo plot of HKU5 S_1_/S_2_ sequences was generated by the Geneious Prime software^126^. Phylogenetic trees of ACE2 were produced using the maximal likelihood method with Q.plant+I+R3 model in IQ-TREE2^127^ (1000 Bootstraps) and rendered with iTOL^128^ (v6).

### Statistical analysis

Most experiments were conducted 2–3 times with 3 biological repeats unless otherwise specified. Representative results were shown. Data were presented by means ± SD as indicated in the figure legends. Unpaired two-tailed t-tests were conducted for all statistical analyses using GraphPad Prism 8. P < 0.05 was considered significant. \**P* < 0.05,\*\**P* < 0.01, \*\*\**P* < 0.005, and \*\*\*\**P* < 0.001.

---

**Figure S1.**
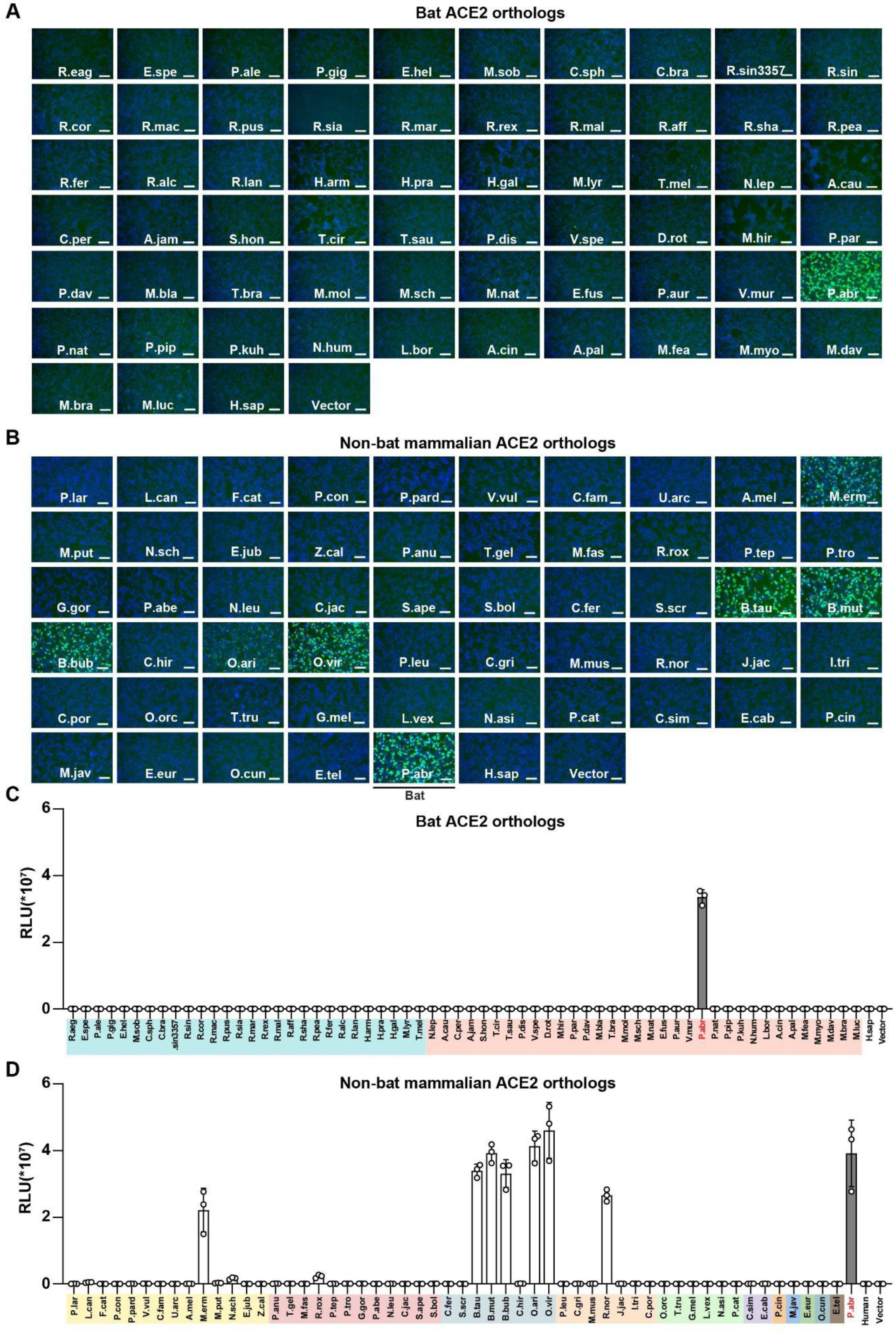
HKU5 utilizes several mammalian ACE2s as receptors. **(A-D)** Binding of the HKU5-19s RBD-hFc to (A-B) and entry of HKU5-19s S VSV pseudovirus into (C-D) HEK293T cells transiently transfected with the indicated bat (A, C) or non-bat (B, D) mammalian ACE2 orthologs. Scale bars: 100 μm. Data are shown as the MEAN ± SD for C-D. n=3 biological replicates. Data representative of two independent experiments for A-B, and a single experiment for C-D.

**Figure S2.**
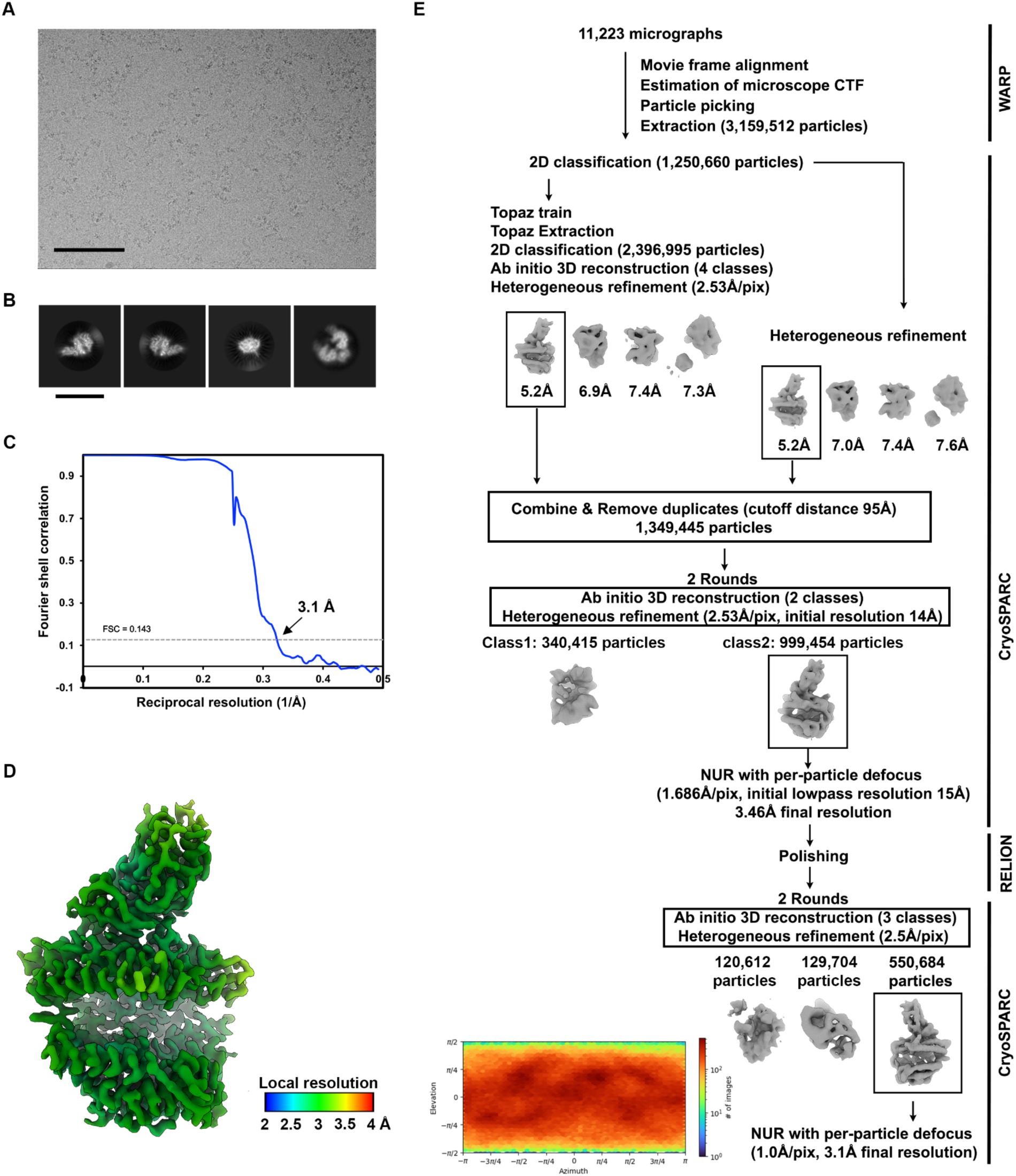
CryoEM data processing of the P.abr ACE2-bound HKU5 RBD dataset. **(A-B)** Representative electron micrograph and 2D class averages of the complex embedded in vitreous ice. Scale bars: 100 nm (A) and 150 Å (B). **(C)** Gold-standard Fourier shell correlation curve. The 0.143 cutoff is indicated by a horizontal dashed line. **(D)** Local resolution estimation calculated using cryoSPARC and plotted on the sharpened map. **(E)** Data processing flowchart. CTF: contrast transfer function; NUR: non-uniform refinement. The angular distribution of particle images calculated using cryoSPARC is shown as a heat map.

**Figure. S3.**
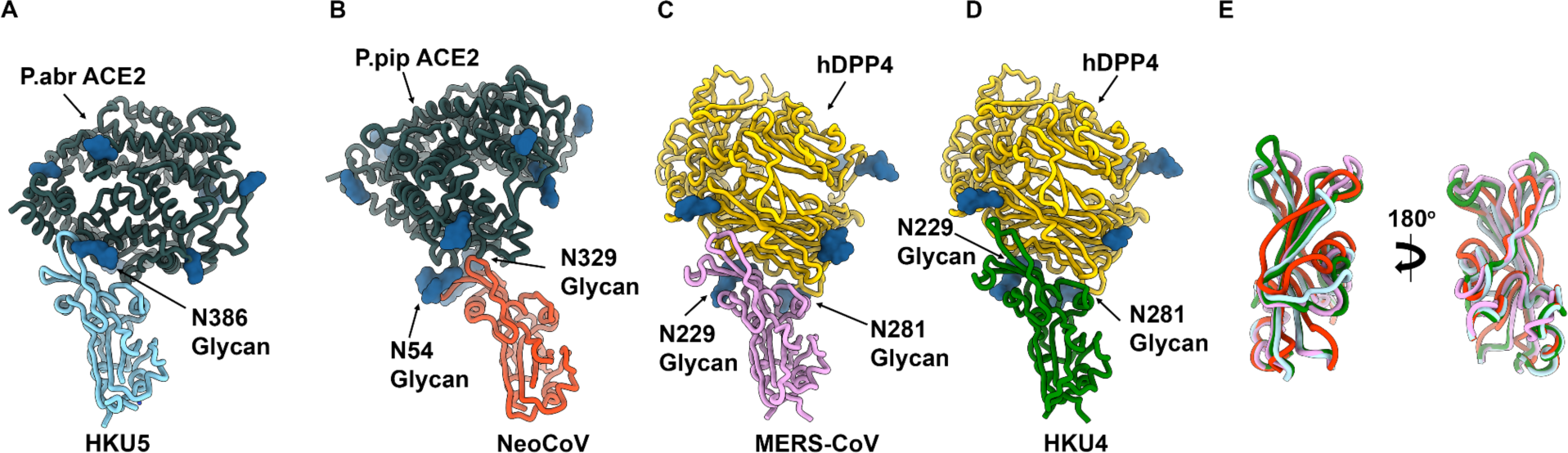
Structural comparisons of the receptor binding domain of HKU5 with HKU4, NeoCoV and MERS-CoV. **(A-D)** Comparison of the structures of the P.abr ACE2-bound HKU5 RBD (PDB 9D32, this study), the P.pip ACE2-bound NeoCoV RBD (PDB 7WPO^39^), hDDP4-bound MERS-CoV RBD (PDB 4KR0^48^) and hDDP4-bound HKU4 RBD (PDB 4QZV^38^). **(E)** Superimposition of the HKU5 (cyan), NeoCoV (orange-red), MERS-CoV (pink) and HKU4 (green) RBMs.

**Figure S4.**
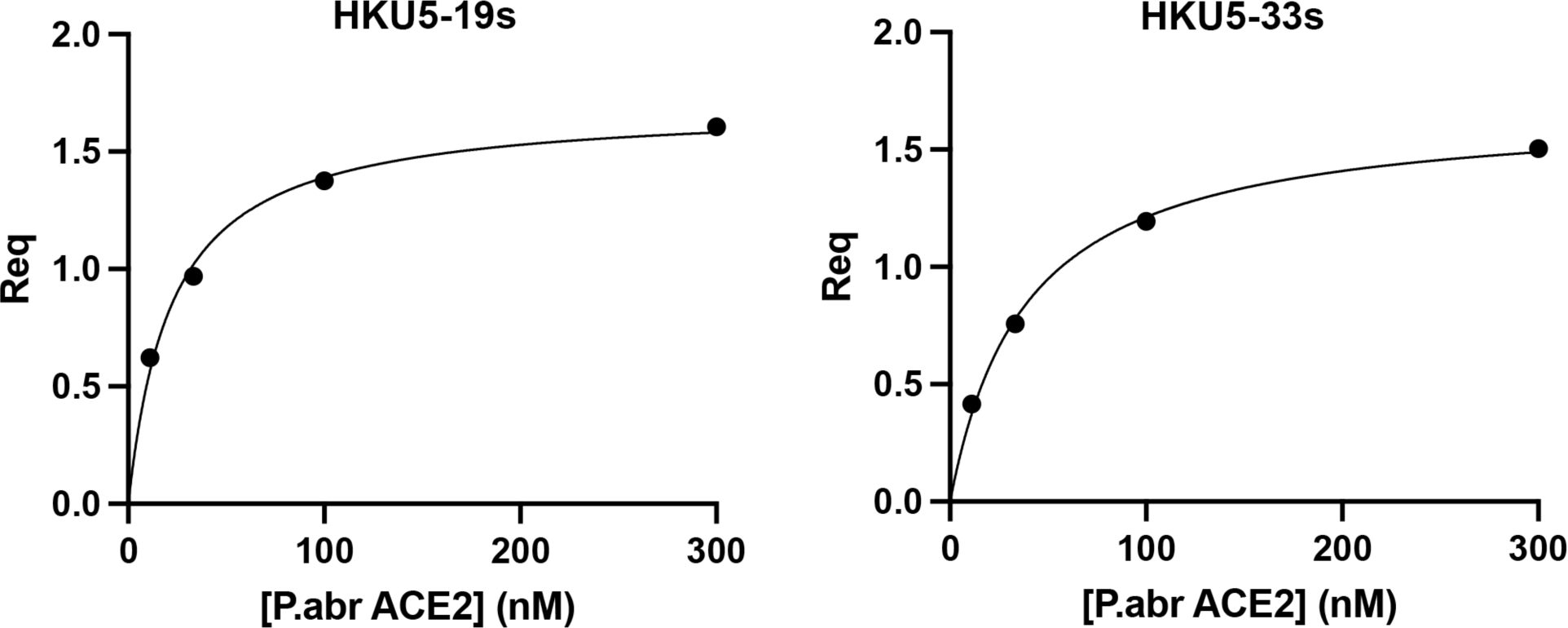
Biolayer interferometry analysis of the P.abr ACE2 ectodomain binding to the immobilized HKU5-19s and HKU5-33s RBDs. Binding avidities were determined by steady state kinetics and are reported as apparent affinities (K_D_, app). One representative out of two technical replicates is shown

**Fig S5.**
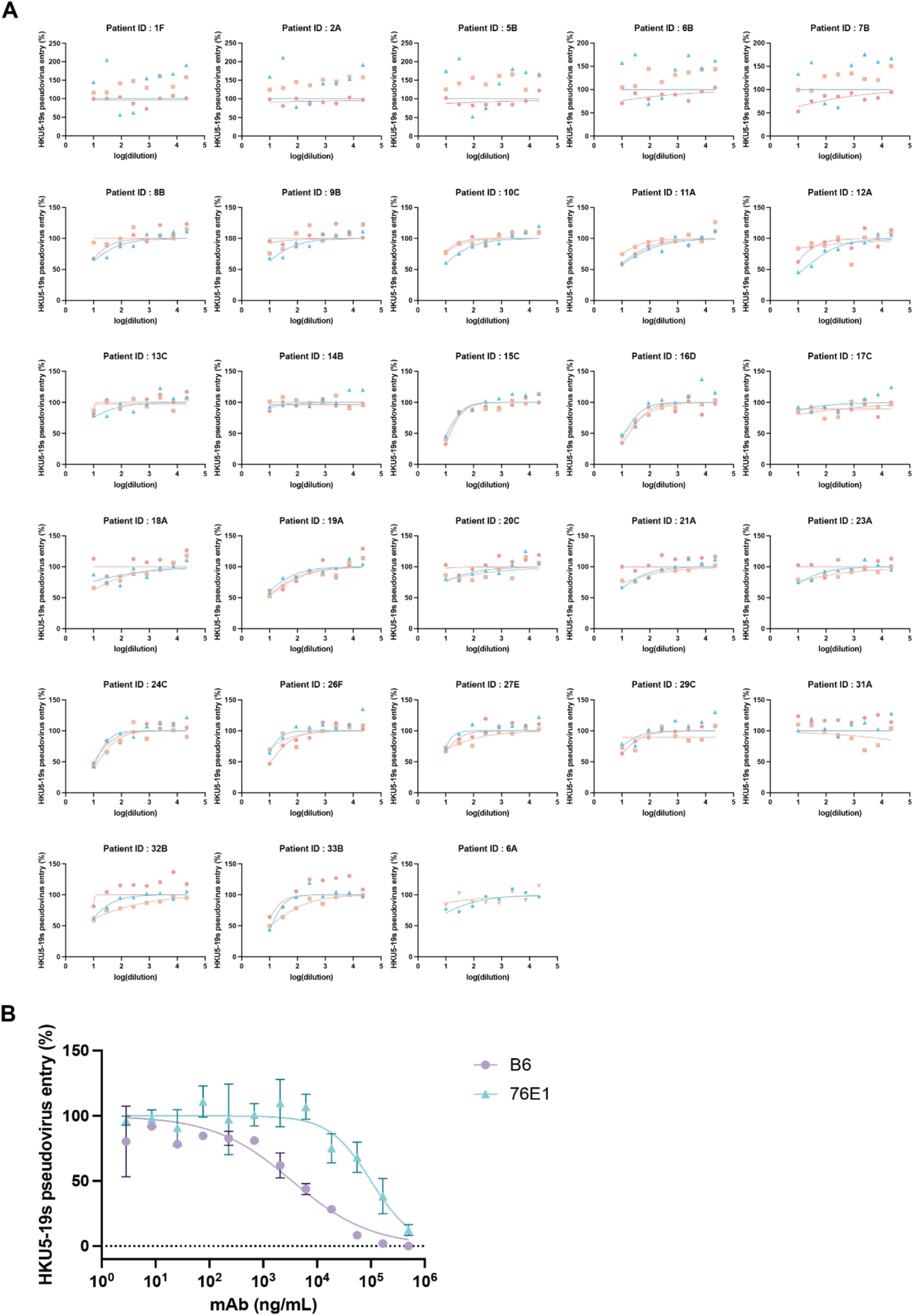
Dose-response curves of HKU5-19s VSV pseudovirus neutralization by MERS-CoV infection-elicited plasma or monoclonal antibodies. **A**, Dose-response curves of HKU5-19s VSV pseudovirus neutralization by MERS-CoV infection-elicited plasma. Each data point represents the mean of two technical replicates and three biological replicates with three distinct batches of pseudoviruses are shown with different colors. **B**, Dose-response curves of HKU5-19s VSV pseudovirus neutralization by fusion machinery-directed monoclonal antibodies. Each data point represents the mean of two technical replicates. A representative curve from three biological replicates is shown.

**Figure S6.**
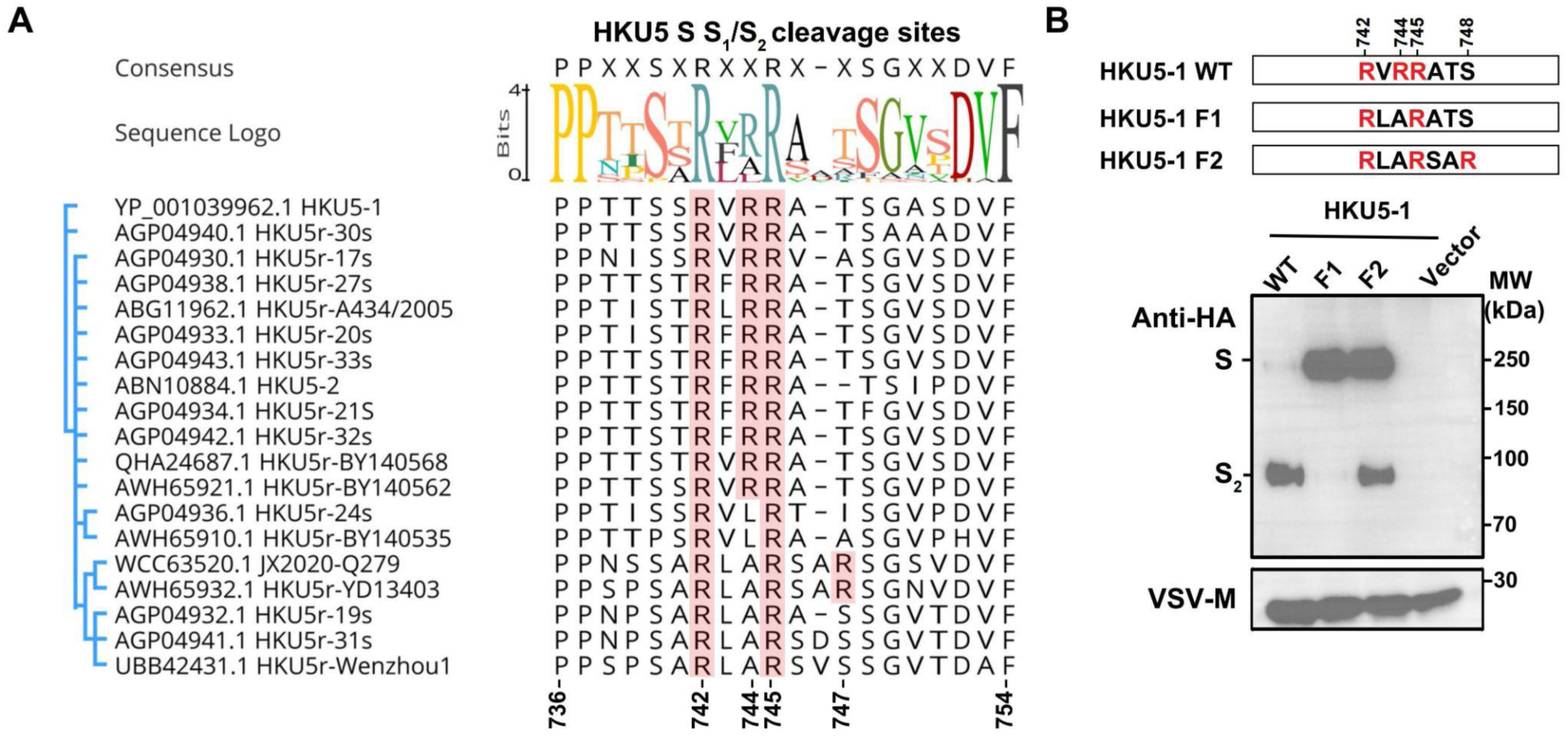
HKU5 S proteolytic processing during biogenesis. **A**, Sequence analysis of S glycoprotein S_1_/S_2_ junction from the indicated HKU5 isolates, with arginine (R) residues highlighted with red background. **B**, Quantification of S glycoprotein proteolytic processing and incorporation in VSV pseudoviruses of wildtype and S_1_/S_2_ HKU5-1 mutants analyzed by Western blot detecting the C-terminal-fused HA tag. VSV-M was used as a loading control. HKU5-1 S residue numbering is shown.

## References

1. Drosten, C., Gunther, S., Preiser, W., van der Werf, S., Brodt, H.R., Becker, S., Rabenau, H., Panning, M., Kolesnikova, L., Fouchier, R.A., et al. (2003). Identification of a novel coronavirus in patients with severe acute respiratory syndrome. N. Engl. J. Med. 348, 1967–1976.

2. Ksiazek, T.G., Erdman, D., Goldsmith, C.S., Zaki, S.R., Peret, T., Emery, S., Tong, S., Urbani, C., Comer, J.A., Lim, W., et al. (2003). A novel coronavirus associated with severe acute respiratory syndrome. N. Engl. J. Med. 348, 1953–1966.

3. Zaki, A.M., van Boheemen, S., Bestebroer, T.M., Osterhaus, A.D., and Fouchier, R.A. (2012). Isolation of a novel coronavirus from a man with pneumonia in Saudi Arabia. N. Engl. J. Med. 367, 1814–1820.

4. Mok, C.K.P., Zhu, A., Zhao, J., Lau, E.H.Y., Wang, J., Chen, Z., Zhuang, Z., Wang, Y., Alshukairi, A.N., Baharoon, S.A., et al. (2020). T-cell responses to MERS coronavirus infection in people with occupational exposure to dromedary camels in Nigeria: an observational cohort study. Lancet Infect. Dis. 10.1016/S1473-3099(20)30599-5.

5. Ngere, I., Hunsperger, E.A., Tong, S., Oyugi, J., Jaoko, W., Harcourt, J.L., Thornburg, N.J., Oyas, H., Muturi, M., Osoro, E.M., et al. (2022). Outbreak of Middle East Respiratory Syndrome Coronavirus in Camels and Probable Spillover Infection to Humans in Kenya. Viruses 14. 10.3390/v14081743.

6. Zhu, N., Zhang, D., Wang, W., Li, X., Yang, B., Song, J., Zhao, X., Huang, B., Shi, W., Lu, R., et al. (2020). A Novel Coronavirus from Patients with Pneumonia in China, 2019. N. Engl. J. Med. 10.1056/NEJMoa2001017.

7. Huang, C., Wang, Y., Li, X., Ren, L., Zhao, J., Hu, Y., Zhang, L., Fan, G., Xu, J., Gu, X., et al. (2020). Clinical features of patients infected with 2019 novel coronavirus in Wuhan, China. Lancet. 10.1016/S0140-6736(20)30183-5.

8. Cameroni, E., Bowen, J.E., Rosen, L.E., Saliba, C., Zepeda, S.K., Culap, K., Pinto, D., VanBlargan, L.A., De Marco, A., di Iulio, J., et al. (2022). Broadly neutralizing antibodies overcome SARS-CoV-2 Omicron antigenic shift. Nature 602, 664–670.

9. Viana, R., Moyo, S., Amoako, D.G., Tegally, H., Scheepers, C., Althaus, C.L., Anyaneji, U.J., Bester, P.A., Boni, M.F., Chand, M., et al. (2022). Rapid epidemic expansion of the SARS-CoV-2 Omicron variant in southern Africa. Nature. 10.1038/d41586-021-03832-5.

10. Ge, X.-Y., Li, J.-L., Yang, X.-L., Chmura, A.A., Zhu, G., Epstein, J.H., Mazet, J.K., Hu, B., Zhang, W., Peng, C., et al. (2013). Isolation and characterization of a bat SARS-like coronavirus that uses the ACE2 receptor. Nature 503, 535–538.

11. Hu, B., Zeng, L.-P., Yang, X.-L., Ge, X.-Y., Zhang, W., Li, B., Xie, J.-Z., Shen, X.-R., Zhang, Y.-Z., Wang, N., et al. (2017). Discovery of a rich gene pool of bat SARS-related coronaviruses provides new insights into the origin of SARS coronavirus. PLoS Pathog. 13, e1006698.

12. Li, W., Shi, Z., Yu, M., Ren, W., Smith, C., Epstein, J.H., Wang, H., Crameri, G., Hu, Z., Zhang, H., et al. (2005). Bats are natural reservoirs of SARS-like coronaviruses. Science 310, 676–679.

13. Yang, X.L., Hu, B., Wang, B., Wang, M.N., Zhang, Q., Zhang, W., Wu, L.J., Ge, X.Y., Zhang, Y.Z., Daszak, P., et al. (2015). Isolation and Characterization of a Novel Bat Coronavirus Closely Related to the Direct Progenitor of Severe Acute Respiratory Syndrome Coronavirus. J. Virol. 90, 3253–3256.

14. Zhou, P., Yang, X.L., Wang, X.G., Hu, B., Zhang, L., Zhang, W., Si, H.R., Zhu, Y., Li, B., Huang, C.L., et al. (2020). A pneumonia outbreak associated with a new coronavirus of probable bat origin. Nature. 10.1038/s41586-020-2012-7.

15. Wang, M., Yan, M., Xu, H., Liang, W., Kan, B., Zheng, B., Chen, H., Zheng, H., Xu, Y., Zhang, E., et al. (2005). SARS-CoV infection in a restaurant from palm civet. Emerg. Infect. Dis. 11, 1860–1865.

16. Consortium, Chinese SARS Molecular Epidemiology (2004). Molecular evolution of the SARS coronavirus during the course of the SARS epidemic in China. Science 303, 1666– 1669.

17. Kan, B., Wang, M., Jing, H., Xu, H., Jiang, X., Yan, M., Liang, W., Zheng, H., Wan, K., Liu, Q., et al. (2005). Molecular evolution analysis and geographic investigation of severe acute respiratory syndrome coronavirus-like virus in palm civets at an animal market and on farms. J. Virol. 79, 11892–11900.

18. Haagmans, B.L., Al Dhahiry, S.H., Reusken, C.B., Raj, V.S., Galiano, M., Myers, R., Godeke, G.J., Jonges, M., Farag, E., Diab, A., et al. (2014). Middle East respiratory syndrome coronavirus in dromedary camels: an outbreak investigation. Lancet Infect. Dis. 14, 140–145.

19. Sabir, J.S., Lam, T.T., Ahmed, M.M., Li, L., Shen, Y., Abo-Aba, S.E., Qureshi, M.I., Abu-Zeid, M., Zhang, Y., Khiyami, M.A., et al. (2016). Co-circulation of three camel coronavirus species and recombination of MERS-CoVs in Saudi Arabia. Science 351, 81–84.

20. Corman, V.M., Kallies, R., Philipps, H., Göpner, G., Müller, M.A., Eckerle, I., Brünink, S., Drosten, C., and Drexler, J.F. (2014). Characterization of a novel betacoronavirus related to middle East respiratory syndrome coronavirus in European hedgehogs. J. Virol. 88, 717– 724.

21. Lau, S.K.P., Luk, H.K.H., Wong, A.C.P., Fan, R.Y.Y., Lam, C.S.F., Li, K.S.M., Ahmed, S.S., Chow, F.W.N., Cai, J.-P., Zhu, X., et al. (2019). Identification of a Novel Betacoronavirus (Merbecovirus) in Amur Hedgehogs from China. Viruses 11. 10.3390/v11110980.

22. Anthony, S.J., Gilardi, K., Menachery, V.D., Goldstein, T., Ssebide, B., Mbabazi, R., Navarrete-Macias, I., Liang, E., Wells, H., Hicks, A., et al. (2017). Further Evidence for Bats as the Evolutionary Source of Middle East Respiratory Syndrome Coronavirus. MBio 8. 10.1128/mBio.00373-17.

23. Walls, A.C., Tortorici, M.A., Bosch, B.J., Frenz, B., Rottier, P.J.M., DiMaio, F., Rey, F.A., and Veesler, D. (2016). Cryo-electron microscopy structure of a coronavirus spike glycoprotein trimer. Nature 531, 114–117.

24. Tortorici, M.A., and Veesler, D. (2019). Structural insights into coronavirus entry. Adv. Virus Res. 105, 93–116.

25. Walls, A.C., Park, Y.-J., Tortorici, M.A., Wall, A., McGuire, A.T., and Veesler, D. (2020). Structure, Function, and Antigenicity of the SARS-CoV-2 Spike Glycoprotein. Cell 181, 281–292.e6.

26. Piccoli, L., Park, Y.J., Tortorici, M.A., Czudnochowski, N., Walls, A.C., Beltramello, M., Silacci-Fregni, C., Pinto, D., Rosen, L.E., Bowen, J.E., et al. (2020). Mapping Neutralizing and Immunodominant Sites on the SARS-CoV-2 Spike Receptor-Binding Domain by Structure-Guided High-Resolution Serology. Cell 183, 1024–1042.e21.

27. Arunachalam, P.S., Walls, A.C., Golden, N., Atyeo, C., Fischinger, S., Li, C., Aye, P., Navarro, M.J., Lai, L., Edara, V.V., et al. (2021). Adjuvanting a subunit COVID-19 vaccine to induce protective immunity. Nature. 10.1038/s41586-021-03530-2.

28. Corbett, K.S., Nason, M.C., Flach, B., Gagne, M., O’Connell, S., Johnston, T.S., Shah, S.N., Edara, V.V., Floyd, K., Lai, L., et al. (2021). Immune correlates of protection by mRNA-1273 vaccine against SARS-CoV-2 in nonhuman primates. Science. 10.1126/science.abj0299.

29. McCallum, M., De Marco, A., Lempp, F.A., Tortorici, M.A., Pinto, D., Walls, A.C., Beltramello, M., Chen, A., Liu, Z., Zatta, F., et al. (2021). N-terminal domain antigenic mapping reveals a site of vulnerability for SARS-CoV-2. Cell 184, 2332–2347.e16.

30. Bowen, J.E., Park, Y.-J., Stewart, C., Brown, J.T., Sharkey, W.K., Walls, A.C., Joshi, A., Sprouse, K.R., McCallum, M., Tortorici, M.A., et al. (2022). SARS-CoV-2 spike conformation determines plasma neutralizing activity elicited by a wide panel of human vaccines. Sci Immunol 7, eadf1421.

31. Barnes, C.O., Jette, C.A., Abernathy, M.E., Dam, K.-M.A., Esswein, S.R., Gristick, H.B., Malyutin, A.G., Sharaf, N.G., Huey-Tubman, K.E., Lee, Y.E., et al. (2020). SARS-CoV-2 neutralizing antibody structures inform therapeutic strategies. Nature 588, 682–687.

32. Millet, J.K., and Whittaker, G.R. (2015). Host cell proteases: Critical determinants of coronavirus tropism and pathogenesis. Virus Res. 202, 120–134.

33. Walls, A.C., Tortorici, M.A., Snijder, J., Xiong, X., Bosch, B.J., Rey, F.A., and Veesler, D. (2017). Tectonic conformational changes of a coronavirus spike glycoprotein promote membrane fusion. Proc. Natl. Acad. Sci. U. S. A. 114, 11157–11162.

34. Niu, Z., Zhang, Z., Gao, X., Du, P., Lu, J., Yan, B., Wang, C., Zheng, Y., Huang, H., and Sun, Q. (2021). N501Y mutation imparts cross-species transmission of SARS-CoV-2 to mice by enhancing receptor binding. Signal Transduct Target Ther 6, 284.

35. McCallum, M., Czudnochowski, N., Rosen, L.E., Zepeda, S.K., Bowen, J.E., Walls, A.C., Hauser, K., Joshi, A., Stewart, C., Dillen, J.R., et al. (2022). Structural basis of SARS-CoV-2 Omicron immune evasion and receptor engagement. Science, eabn8652.

36. Raj, V.S., Mou, H., Smits, S.L., Dekkers, D.H.W., Müller, M.A., Dijkman, R., Muth, D., Demmers, J.A.A., Zaki, A., Fouchier, R.A.M., et al. (2013). Dipeptidyl peptidase 4 is a functional receptor for the emerging human coronavirus-EMC. Nature 495, 251–254.

37. Chen, J., Yang, X., Si, H., Gong, Q., Que, T., Li, J., Li, Y., Wu, C., Zhang, W., Chen, Y., et al. (2023). A bat MERS-like coronavirus circulates in pangolins and utilizes human DPP4 and host proteases for cell entry. Cell 186, 850–863.e16.

38. Wang, Q., Qi, J., Yuan, Y., Xuan, Y., Han, P., Wan, Y., Ji, W., Li, Y., Wu, Y., Wang, J., et al. (2014). Bat origins of MERS-CoV supported by bat coronavirus HKU4 usage of human receptor CD26. Cell Host Microbe 16, 328–337.

39. Xiong, Q., Cao, L., Ma, C., Tortorici, M.A., Liu, C., Si, J., Liu, P., Gu, M., Walls, A.C., Wang, C., et al. (2022). Close relatives of MERS-CoV in bats use ACE2 as their functional receptors. Nature 612, 748–757.

40. Ma, C., Liu, C., Xiong, Q., Gu, M., Shi, L., Wang, C., Si, J., Tong, F., Liu, P., Huang, M., et al. (2023). Broad host tropism of ACE2-using MERS-related coronaviruses and determinants restricting viral recognition. Cell Discov 9, 57.

41. Woo, P.C., Lau, S.K., Li, K.S., Poon, R.W., Wong, B.H., Tsoi, H.W., Yip, B.C., Huang, Y., Chan, K.H., and Yuen, K.Y. (2006). Molecular diversity of coronaviruses in bats. Virology 351, 180–187.

42. Ithete, N.L., Stoffberg, S., Corman, V.M., Cottontail, V.M., Richards, L.R., Schoeman, M.C., Drosten, C., Drexler, J.F., and Preiser, W. (2013). Close relative of human Middle East respiratory syndrome coronavirus in bat, South Africa. Emerg. Infect. Dis. 19, 1697–1699.

43. Yan, H., Jiao, H., Liu, Q., Zhang, Z., Xiong, Q., Wang, B.-J., Wang, X., Guo, M., Wang, L.-F., Lan, K., et al. (2021). ACE2 receptor usage reveals variation in susceptibility to SARS-CoV and SARS-CoV-2 infection among bat species. Nat Ecol Evol 5, 600–608.

44. Ma, C.-B., Liu, C., Park, Y.-J., Tang, J., Chen, J., Xiong, Q., Lee, J., Stewart, C., Asarnow, D., Brown, J., et al. (2024). Multiple independent acquisitions of ACE2 usage in MERS-related coronaviruses. bioRxiv, 2023.10.02.560486. 10.1101/2023.10.02.560486.

45. Liu, P., Huang, M.-L., Guo, H., Si, J.-Y., Chen, Y.-M., Wang, C.-L., Yu, X., Shi, L.-L., Xiong, Q., Ma, C.-B., et al. (2024). Engineering customized viral receptors for various coronaviruses. bioRxiv, 2024.03.03.583237. 10.1101/2024.03.03.583237.

46. Menachery, V.D., Dinnon, K.H., 3rd, Yount, B.L., Jr, McAnarney, E.T., Gralinski, L.E., Hale, A., Graham, R.L., Scobey, T., Anthony, S.J., Wang, L., et al. (2020). Trypsin Treatment Unlocks Barrier for Zoonotic Bat Coronavirus Infection. J. Virol. 94. 10.1128/JVI.01774-19.

47. Letko, M. (2024). Functional assessment of cell entry and receptor use for merbecoviruses. bioRxiv, 2024.03.13.584892. 10.1101/2024.03.13.584892.

48. Lu, G., Hu, Y., Wang, Q., Qi, J., Gao, F., Li, Y., Zhang, Y., Zhang, W., Yuan, Y., Bao, J., et al. (2013). Molecular basis of binding between novel human coronavirus MERS-CoV and its receptor CD26. Nature 500, 227–231.

49. Wang, N., Shi, X., Jiang, L., Zhang, S., Wang, D., Tong, P., Guo, D., Fu, L., Cui, Y., Liu, X., et al. (2013). Structure of MERS-CoV spike receptor-binding domain complexed with human receptor DPP4. Cell Res. 23, 986–993.

50. Addetia, A., Piccoli, L., Case, J.B., Park, Y.-J., Beltramello, M., Guarino, B., Dang, H., de Melo, G.D., Pinto, D., Sprouse, K., et al. (2023). Neutralization, effector function and immune imprinting of Omicron variants. Nature 621, 592–601.

51. Starr, T.N., Greaney, A.J., Hannon, W.W., Loes, A.N., Hauser, K., Dillen, J.R., Ferri, E., Farrell, A.G., Dadonaite, B., McCallum, M., et al. (2022). Shifting mutational constraints in the SARS-CoV-2 receptor-binding domain during viral evolution. Preprint, 10.1126/science.abo7896.

52. Starr, T.N., Greaney, A.J., Hilton, S.K., Ellis, D., Crawford, K.H.D., Dingens, A.S., Navarro, M.J., Bowen, J.E., Tortorici, M.A., Walls, A.C., et al. (2020). Deep Mutational Scanning of SARS-CoV-2 Receptor Binding Domain Reveals Constraints on Folding and ACE2 Binding. Cell 182, 1295–1310.e20.

53. Wang, Q., Guo, Y., Liu, L., Schwanz, L.T., Li, Z., Nair, M.S., Ho, J., Zhang, R.M., Iketani, S., Yu, J., et al. (2023). Antigenicity and receptor affinity of SARS-CoV-2 BA.2.86 spike. Nature. 10.1038/s41586-023-06750-w.

54. Yue, C., Song, W., Wang, L., Jian, F., Chen, X., Gao, F., Shen, Z., Wang, Y., Wang, X., and Cao, Y. (2023). ACE2 binding and antibody evasion in enhanced transmissibility of XBB.1.5. Lancet Infect. Dis. 10.1016/S1473-3099(23)00010-5.

55. McCallum, M., Walls, A.C., Sprouse, K.R., Bowen, J.E., Rosen, L.E., Dang, H.V., De Marco, A., Franko, N., Tilles, S.W., Logue, J., et al. (2021). Molecular basis of immune evasion by the Delta and Kappa SARS-CoV-2 variants. Science, eabl8506.

56. Bowen, J.E., Addetia, A., Dang, H.V., Stewart, C., Brown, J.T., Sharkey, W.K., Sprouse, K.R., Walls, A.C., Mazzitelli, I.G., Logue, J.K., et al. (2022). Omicron spike function and neutralizing activity elicited by a comprehensive panel of vaccines. Science 377, 890–894.

57. Lan, J., Ge, J., Yu, J., Shan, S., Zhou, H., Fan, S., Zhang, Q., Shi, X., Wang, Q., Zhang, L., et al. (2020). Structure of the SARS-CoV-2 spike receptor-binding domain bound to the ACE2 receptor. Nature. 10.1038/s41586-020-2180-5.

58. Addetia, A., Stewart, C., Seo, A.J., Sprouse, K.R., Asiri, A.Y., Al-Mozaini, M., Memish, Z.A., Alshukairi, A.N., and Veesler, D. (2024). Mapping immunodominant sites on the MERS-CoV spike glycoprotein targeted by infection-elicited antibodies in humans. Cell Rep. 43, 114530.

59. Sauer, M.M., Tortorici, M.A., Park, Y.-J., Walls, A.C., Homad, L., Acton, O.J., Bowen, J.E., Wang, C., Xiong, X., de van der Schueren, W., et al. (2021). Structural basis for broad coronavirus neutralization. Nat. Struct. Mol. Biol. 28, 478–486.

60. Pinto, D., Sauer, M.M., Czudnochowski, N., Low, J.S., Alejandra Tortorici, M., Housley, M.P., Noack, J., Walls, A.C., Bowen, J.E., Guarino, B., et al. (2021). Broad betacoronavirus neutralization by a stem helix–specific human antibody. Science. 10.1126/science.abj3321.

61. Sun, X., Yi, C., Zhu, Y., Ding, L., Xia, S., Chen, X., Liu, M., Gu, C., Lu, X., Fu, Y., et al. (2022). Neutralization mechanism of a human antibody with pan-coronavirus reactivity including SARS-CoV-2. Nat Microbiol 7, 1063–1074.

62. Low, J.S., Jerak, J., Tortorici, M.A., McCallum, M., Pinto, D., Cassotta, A., Foglierini, M., Mele, F., Abdelnabi, R., Weynand, B., et al. (2022). ACE2-binding exposes the SARS-CoV-2 fusion peptide to broadly neutralizing coronavirus antibodies. Science, eabq2679.

63. Hoffmann, M., Kleine-Weber, H., Schroeder, S., Krüger, N., Herrler, T., Erichsen, S., Schiergens, T.S., Herrler, G., Wu, N.H., Nitsche, A., et al. (2020). SARS-CoV-2 Cell Entry Depends on ACE2 and TMPRSS2 and Is Blocked by a Clinically Proven Protease Inhibitor. Cell 181, 271–280.e8.

64. Fraser, B.J., Beldar, S., Seitova, A., Hutchinson, A., Mannar, D., Li, Y., Kwon, D., Tan, R., Wilson, R.P., Leopold, K., et al. (2022). Structure and activity of human TMPRSS2 protease implicated in SARS-CoV-2 activation. Nat. Chem. Biol. 18, 963–971.

65. Icho, S., Rujas, E., Muthuraman, K., Tam, J., Liang, H., Landreth, S., Liao, M., Falzarano, D., Julien, J.-P., and Melnyk, R.A. (2022). Dual Inhibition of Vacuolar-ATPase and TMPRSS2 Is Required for Complete Blockade of SARS-CoV-2 Entry into Cells. Antimicrob. Agents Chemother. 66, e0043922.

66. Park, J.-E., Li, K., Barlan, A., Fehr, A.R., Perlman, S., McCray, P.B., Jr, and Gallagher, T. (2016). Proteolytic processing of Middle East respiratory syndrome coronavirus spikes expands virus tropism. Proc. Natl. Acad. Sci. U. S. A. 113, 12262–12267.

67. Millet, J.K., and Whittaker, G.R. (2014). Host cell entry of Middle East respiratory syndrome coronavirus after two-step, furin-mediated activation of the spike protein. Proc. Natl. Acad. Sci. U. S. A. 111, 15214–15219.

68. Biniossek, M.L., Nägler, D.K., Becker-Pauly, C., and Schilling, O. (2011). Proteomic identification of protease cleavage sites characterizes prime and non-prime specificity of cysteine cathepsins B, L, and S. J. Proteome Res. 10, 5363–5373.

69. Yang, Y., Liu, C., Du, L., Jiang, S., Shi, Z., Baric, R.S., and Li, F. (2015). Two Mutations Were Critical for Bat-to-Human Transmission of Middle East Respiratory Syndrome Coronavirus. J. Virol. 89, 9119–9123.

70. Xia, S., Yan, L., Xu, W., Agrawal, A.S., Algaissi, A., Tseng, C.-T.K., Wang, Q., Du, L., Tan, W., Wilson, I.A., et al. (2019). A pan-coronavirus fusion inhibitor targeting the HR1 domain of human coronavirus spike. Sci Adv 5, eaav4580.

71. Xia, S., Liu, M., Wang, C., Xu, W., Lan, Q., Feng, S., Qi, F., Bao, L., Du, L., Liu, S., et al. (2020). Inhibition of SARS-CoV-2 (previously 2019-nCoV) infection by a highly potent pan-coronavirus fusion inhibitor targeting its spike protein that harbors a high capacity to mediate membrane fusion. Cell Res. 30, 343–355.

72. de Wit, E., Feldmann, F., Cronin, J., Jordan, R., Okumura, A., Thomas, T., Scott, D., Cihlar, T., and Feldmann, H. (2020). Prophylactic and therapeutic remdesivir (GS-5734) treatment in the rhesus macaque model of MERS-CoV infection. Proc. Natl. Acad. Sci. U. S. A. 117, 6771–6776.

73. Sheahan, T.P., Sims, A.C., Leist, S.R., Schäfer, A., Won, J., Brown, A.J., Montgomery, S.A., Hogg, A., Babusis, D., Clarke, M.O., et al. (2020). Comparative therapeutic efficacy of remdesivir and combination lopinavir, ritonavir, and interferon beta against MERS-CoV. Nat. Commun. 11, 222.

74. Kokic, G., Hillen, H.S., Tegunov, D., Dienemann, C., Seitz, F., Schmitzova, J., Farnung, L., Siewert, A., Höbartner, C., and Cramer, P. (2021). Mechanism of SARS-CoV-2 polymerase stalling by remdesivir. Nat. Commun. 12, 279.

75. Wu, K., Li, W., Peng, G., and Li, F. (2009). Crystal structure of NL63 respiratory coronavirus receptor-binding domain complexed with its human receptor. Proc. Natl. Acad. Sci. U. S. A. 106, 19970–19974.

76. Yan, R., Zhang, Y., Li, Y., Xia, L., Guo, Y., and Zhou, Q. (2020). Structural basis for the recognition of SARS-CoV-2 by full-length human ACE2. Science 367, 1444–1448.

77. Yang, Y., Du, L., Liu, C., Wang, L., Ma, C., Tang, J., Baric, R.S., Jiang, S., and Li, F. (2014). Receptor usage and cell entry of bat coronavirus HKU4 provide insight into bat-to-human transmission of MERS coronavirus. Proc. Natl. Acad. Sci. U. S. A. 111, 12516– 12521.

78. Li, F., Li, W., Farzan, M., and Harrison, S.C. (2005). Structure of SARS coronavirus spike receptor-binding domain complexed with receptor. Science 309, 1864–1868.

79. Reguera, J., Santiago, C., Mudgal, G., Ordono, D., Enjuanes, L., and Casasnovas, J.M. (2012). Structural bases of coronavirus attachment to host aminopeptidase N and its inhibition by neutralizing antibodies. PLoS Pathog. 8, e1002859.

80. Tortorici, M.A., Walls, A.C., Joshi, A., Park, Y.-J., Eguia, R.T., Miranda, M.C., Kepl, E., Dosey, A., Stevens-Ayers, T., Boeckh, M.J., et al. (2022). Structure, receptor recognition, and antigenicity of the human coronavirus CCoV-HuPn-2018 spike glycoprotein. Cell 185, 2279–2291.e17.

81. Ji, W., Peng, Q., Fang, X., Li, Z., Li, Y., Xu, C., Zhao, S., Li, J., Chen, R., Mo, G., et al. (2022). Structures of a deltacoronavirus spike protein bound to porcine and human receptors. Nat. Commun. 13, 1467.

82. Wong, A.H., and Rini, J.M. (2017). Crystal structure of the human coronavirus 229E spike protein receptor binding domain in complex with human aminopeptidase N. Preprint at Worldwide Protein Data Bank, 10.2210/pdb6atk/pdb.

83. Greaney, A.J., Loes, A.N., Gentles, L.E., Crawford, K.H.D., Starr, T.N., Malone, K.D., Chu, H.Y., and Bloom, J.D. (2021). Antibodies elicited by mRNA-1273 vaccination bind more broadly to the receptor binding domain than do those from SARS-CoV-2 infection. Sci. Transl. Med. 13. 10.1126/scitranslmed.abi9915.

84. Lee, J., Zepeda, S.K., Park, Y.-J., Taylor, A.L., Quispe, J., Stewart, C., Leaf, E.M., Treichel, C., Corti, D., King, N.P., et al. (2023). Broad receptor tropism and immunogenicity of a clade 3 sarbecovirus. Cell Host Microbe 31, 1961–1973.e11.

85. Walls, A.C., Miranda, M.C., Schäfer, A., Pham, M.N., Greaney, A., Arunachalam, P.S., Navarro, M.-J., Tortorici, M.A., Rogers, K., O’Connor, M.A., et al. (2021). Elicitation of broadly protective sarbecovirus immunity by receptor-binding domain nanoparticle vaccines. Cell. 10.1016/j.cell.2021.09.015.

86. Chao, C.W., Sprouse, K.R., Miranda, M.C., Catanzaro, N.J., Hubbard, M.L., Addetia, A., Stewart, C., Brown, J.T., Dosey, A., Valdez, A., et al. (2024). Protein nanoparticle vaccines induce potent neutralizing antibody responses against MERS-CoV. bioRxiv. 10.1101/2024.03.13.584735.

87. Song, G., He, W.-T., Callaghan, S., Anzanello, F., Huang, D., Ricketts, J., Torres, J.L., Beutler, N., Peng, L., Vargas, S., et al. (2021). Cross-reactive serum and memory B-cell responses to spike protein in SARS-CoV-2 and endemic coronavirus infection. Nat. Commun. 12, 2938.

88. Zhou, P., Song, G., Liu, H., Yuan, M., He, W.-T., Beutler, N., Zhu, X., Tse, L.V., Martinez, D.R., Schäfer, A., et al. (2023). Broadly neutralizing anti-S2 antibodies protect against all three human betacoronaviruses that cause deadly disease. Immunity 56, 669–686.e7.

89. Tortorici, M.A., Czudnochowski, N., Starr, T.N., Marzi, R., Walls, A.C., Zatta, F., Bowen, J.E., Jaconi, S., Di Iulio, J., Wang, Z., et al. (2021). Broad sarbecovirus neutralization by a human monoclonal antibody. Nature. 10.1038/s41586-021-03817-4.

90. Starr, T.N., Czudnochowski, N., Liu, Z., Zatta, F., Park, Y.-J., Addetia, A., Pinto, D., Beltramello, M., Hernandez, P., Greaney, A.J., et al. (2021). SARS-CoV-2 RBD antibodies that maximize breadth and resistance to escape. Nature. 10.1038/s41586-021-03807-6.

91. Schwegmann-Weßels, C., Glende, J., Ren, X., Qu, X., Deng, H., Enjuanes, L., and Herrler, G. (2009). Comparison of vesicular stomatitis virus pseudotyped with the S proteins from a porcine and a human coronavirus. J. Gen. Virol. 90, 1724–1729.

92. Reed, L.J., and Muench, H. (1938). A Simple Method for Estimating Fifty Per Cent Endpoints (Lancaster Press, Incorporated).

93. Russo, C.J., and Passmore, L.A. (2014). Electron microscopy: Ultrastable gold substrates for electron cryomicroscopy. Science 346, 1377–1380.

94. Suloway, C., Pulokas, J., Fellmann, D., Cheng, A., Guerra, F., Quispe, J., Stagg, S., Potter, C.S., and Carragher, B. (2005). Automated molecular microscopy: the new Leginon system. J. Struct. Biol. 151, 41–60.

95. Tegunov, D., and Cramer, P. (2019). Real-time cryo-electron microscopy data preprocessing with Warp. Nat. Methods 16, 1146–1152.

96. Punjani, A., Rubinstein, J.L., Fleet, D.J., and Brubaker, M.A. (2017). cryoSPARC: algorithms for rapid unsupervised cryo-EM structure determination. Nat. Methods 14, 290– 296.

97. Bepler, T., Kelley, K., Noble, A.J., and Berger, B. (2020). Topaz-Denoise: general deep denoising models for cryoEM and cryoET. Nat. Commun. 11, 5208.

98. Punjani, A., Zhang, H., and Fleet, D.J. (2020). Non-uniform refinement: adaptive regularization improves single-particle cryo-EM reconstruction. Nat. Methods 17, 1214– 1221.

99. Zivanov, J., Nakane, T., Forsberg, B.O., Kimanius, D., Hagen, W.J., Lindahl, E., and Scheres, S.H. (2018). New tools for automated high-resolution cryo-EM structure determination in RELION-3. Elife 7. 10.7554/eLife.42166.

100. Asarnow, D., Palovcak, E., and Cheng, Y. (2019). UCSF pyem v0. 5. Zenodo 10.5281/zenodo3576630, 2019.

101. Zivanov, J., Nakane, T., and Scheres, S.H.W. (2019). A Bayesian approach to beam-induced motion correction in cryo-EM single-particle analysis. IUCrJ 6, 5–17.

102. Rosenthal, P.B., and Henderson, R. (2003). Optimal determination of particle orientation, absolute hand, and contrast loss in single-particle electron cryomicroscopy. J. Mol. Biol. 333, 721–745.

103. Chen, S., McMullan, G., Faruqi, A.R., Murshudov, G.N., Short, J.M., Scheres, S.H., and Henderson, R. (2013). High-resolution noise substitution to measure overfitting and validate resolution in 3D structure determination by single particle electron cryomicroscopy. Ultramicroscopy 135, 24–35.

104. Pettersen, E.F., Goddard, T.D., Huang, C.C., Couch, G.S., Greenblatt, D.M., Meng, E.C., and Ferrin, T.E. (2004). UCSF Chimera--a visualization system for exploratory research and analysis. J. Comput. Chem. 25, 1605–1612.

105. Emsley, P., Lohkamp, B., Scott, W.G., and Cowtan, K. (2010). Features and development of Coot. Acta Crystallogr. D Biol. Crystallogr. 66, 486–501.

106. Abramson, J., Adler, J., Dunger, J., Evans, R., Green, T., Pritzel, A., Ronneberger, O., Willmore, L., Ballard, A.J., Bambrick, J., et al. (2024). Accurate structure prediction of biomolecular interactions with AlphaFold 3. Nature 630, 493–500.

107. Frenz, B., Rämisch, S., Borst, A.J., Walls, A.C., Adolf-Bryfogle, J., Schief, W.R., Veesler, D., and DiMaio, F. (2019). Automatically Fixing Errors in Glycoprotein Structures with Rosetta. Structure 27, 134–139.e3.

108. Wang, R.Y., Song, Y., Barad, B.A., Cheng, Y., Fraser, J.S., and DiMaio, F. (2016). Automated structure refinement of macromolecular assemblies from cryo-EM maps using Rosetta. Elife 5. 10.7554/eLife.17219.

109. Liebschner, D., Afonine, P.V., Baker, M.L., Bunkóczi, G., Chen, V.B., Croll, T.I., Hintze, B., Hung, L.W., Jain, S., McCoy, A.J., et al. (2019). Macromolecular structure determination using X-rays, neutrons and electrons: recent developments in Phenix. Acta Crystallogr D Struct Biol 75, 861–877.

110. Chen, V.B., Arendall, W.B., Headd, J.J., Keedy, D.A., Immormino, R.M., Kapral, G.J., Murray, L.W., Richardson, J.S., and Richardson, D.C. (2010). MolProbity: all-atom structure validation for macromolecular crystallography. Acta Crystallogr. D Biol. Crystallogr. 66, 12– 21.

111. Barad, B.A., Echols, N., Wang, R.Y., Cheng, Y., DiMaio, F., Adams, P.D., and Fraser, J.S. (2015). EMRinger: side chain-directed model and map validation for 3D cryo-electron microscopy. Nat. Methods 12, 943–946.

112. Agirre, J., Iglesias-Fernández, J., Rovira, C., Davies, G.J., Wilson, K.S., and Cowtan, K.D. (2015). Privateer: software for the conformational validation of carbohydrate structures. Nat. Struct. Mol. Biol. 22, 833–834.

113. Stott, C.J., Sawattrakool, K., Saeng-Chuto, K., Tantituvanont, A., and Nilubol, D. (2022). The phylodynamics of emerging porcine deltacoronavirus in Southeast Asia. Transbound. Emerg. Dis. 69, 2816–2827.

114. Ye, X., Chen, Y., Zhu, X., Guo, J., Xie, D., Hou, Z., Xu, S., Zhou, J., Fang, L., Wang, D., et al. (2020). Cross-species transmission of deltacoronavirus and the origin of porcine deltacoronavirus. Evol. Appl. 13, 2246–2253.

115. He, W.-T., Ji, X., He, W., Dellicour, S., Wang, S., Li, G., Zhang, L., Gilbert, M., Zhu, H., Xing, G., et al. (2020). Genomic Epidemiology, Evolution, and Transmission Dynamics of Porcine Deltacoronavirus. Mol. Biol. Evol. 37, 2641–2654.

116. Hsueh, F.-C., Wu, C.-N., Lin, M.Y.-C., Hsu, F.-Y., Lin, C.-F., Chang, H.-W., Lin, J.-H., Liu, H.-F., Chiou, M.-T., Chan, K.R., et al. (2021). Phylodynamic analysis and spike protein mutations in porcine deltacoronavirus with a new variant introduction in Taiwan. Virus Evol 7, veab096.

117. Lednicky, J.A., Tagliamonte, M.S., White, S.K., Elbadry, M.A., Alam, M.M., Stephenson, C.J., Bonny, T.S., Loeb, J.C., Telisma, T., Chavannes, S., et al. (2021). Independent infections of porcine deltacoronavirus among Haitian children. Nature 600, 133–137.

118. Fu, L., Niu, B., Zhu, Z., Wu, S., and Li, W. (2012). CD-HIT: accelerated for clustering the next-generation sequencing data. Bioinformatics 28, 3150–3152.

119. Rozewicki, J., Li, S., Amada, K.M., Standley, D.M., and Katoh, K. (2019). MAFFT-DASH: integrated protein sequence and structural alignment. Nucleic Acids Res. 47, W5–W10.

120. Kosakovsky Pond, S.L., Posada, D., Gravenor, M.B., Woelk, C.H., and Frost, S.D.W. (2006). GARD: a genetic algorithm for recombination detection. Bioinformatics 22, 3096– 3098.

121. Stamatakis, A. (2014). RAxML version 8: a tool for phylogenetic analysis and post-analysis of large phylogenies. Bioinformatics 30, 1312–1313.

122. Tolentino, J.E., Lytras, S., Ito, J., and Sato, K. (2024). Recombination analysis on the receptor switching event of MERS-CoV and its close relatives: implications for the emergence of MERS-CoV. Virol. J. 21, 84.

123. Tan, C.C.S., Trew, J., Peacock, T.P., Mok, K.Y., Hart, C., Lau, K., Ni, D., Orme, C.D.L., Ransome, E., Pearse, W.D., et al. (2023). Genomic screening of 16 UK native bat species through conservationist networks uncovers coronaviruses with zoonotic potential. Nat. Commun. 14, 3322.

124. Edgar, R.C. (2004). MUSCLE: multiple sequence alignment with high accuracy and high throughput. Nucleic Acids Res. 32, 1792–1797.

125. Kumar, S., Stecher, G., Li, M., Knyaz, C., and Tamura, K. (2018). MEGA X: Molecular Evolutionary Genetics Analysis across Computing Platforms. Mol. Biol. Evol. 35, 1547– 1549.

126. Kearse, M., Moir, R., Wilson, A., Stones-Havas, S., Cheung, M., Sturrock, S., Buxton, S., Cooper, A., Markowitz, S., Duran, C., et al. (2012). Geneious Basic: an integrated and extendable desktop software platform for the organization and analysis of sequence data. Bioinformatics 28, 1647–1649.

127. Minh, B.Q., Schmidt, H.A., Chernomor, O., Schrempf, D., Woodhams, M.D., von Haeseler, A., and Lanfear, R. (2020). IQ-TREE 2: New Models and Efficient Methods for Phylogenetic Inference in the Genomic Era. Mol. Biol. Evol. 37, 1530–1534.

128. Letunic, I., and Bork, P. (2024). Interactive Tree of Life (iTOL) v6: recent updates to the phylogenetic tree display and annotation tool. Nucleic Acids Res. 52, W78–W82.

